# Engineering cellular communication between light-activated synthetic cells and bacteria

**DOI:** 10.1101/2022.07.22.500923

**Authors:** Jefferson M. Smith, Denis Hartmann, Michael J. Booth

**Affiliations:** Department of Chemistry, University of Oxford, Mansfield Road, OX1 3TA, Oxford, UK; Department of Chemistry, University College London, 20 Gordon Street, WC1H 0AJ, London UK

## Abstract

Gene-expressing compartments assembled from simple, modular parts, are a versatile platform for creating minimal synthetic cells. In a manner synonymous with natural cells, these life-like assemblies utilise information encoded in DNA within the compartment’s interior to dictate which proteins are expressed and consequently the overall function of the synthetic cell. Hence, by incorporating gene regulatory motifs into the DNA templates, *in situ* gene expression can be controlled according to specific stimuli. In this work, cell-free protein synthesis within synthetic cells was controlled using light by encoding genes of interest on light-activated DNA templates. Light-activated DNA contained a photocleavable blockade within the T7 promoter region that tightly repressed transcription until the blocking groups were removed with UV light. In this way, synthetic cells were activated remotely, in a spatiotemporally controlled manner. By applying this strategy to the expression of an acyl homo-serine lactone synthetase, BjaI, quorum-sensing based communication between synthetic cells and bacteria was controlled with light. This work provides a framework for the remote-controlled production and delivery of small molecules from non-living matter to living matter, with applications in biology and medicine.

## Introduction

Re-programming living systems to adopt novel functions outside their natural capabilities is a notoriously difficult endeavour. One approach that has helped to remedy this involves reducing the system’s complexity and constructing the desired biological process from the bottom-up using simple, modular parts. In this manner, cell-free systems have been exploited to rapidly optimise biosynthetic pathways *in vitro* [1], create portable, low-cost biosensors against a variety of targets [2, 3], and even create minimalistic synthetic cells from scratch [4]. Synthetic cells are non-living, cell-like compartments capable of basic life-like behaviours. While the term synthetic cell is broad and encompasses various compartmentalised systems [5], liposomes functionalised with a cell-free protein synthesis (CFPS) system have become a prevalent model system, offering a neutral breadboard into which biological circuitry can be assembled. Like living cells, these liposome-based synthetic cells are bound by lipid membranes and their behaviours, including differentiation [6] and communication [7–14], are programmed according to DNA templates. However, their makeup is not restricted by the design rules imposed by nature, and they can also accommodate non-biological components alongside their biological components to attain entirely new functionalities.

Despite the seemingly endless array of parts available to engineer synthetic cells, comparatively, few have been explored [15]. For example, of the enumerable gene regulatory networks observed in nature, *in situ* gene expression inside synthetic cells is often controlled at either the DNA and mRNA level using only a few well-defined small molecule-sensitive transcription factors [9, 12, 14] and translational riboswitches [7, 9, 16, 17], respectively. While these tools exhibit stringent control over gene expression *in vivo* [18, 19], this is not reflected in their performance *in vitro*, as gene expression is often leaky and occurs even in the absence of the cognate ligand [7, 16]. A small subset of parts have also been recycled in most synthetic cell communication networks [15]. Synthetic cells typically communicate with other synthetic cells or living cells by releasing entrapped membrane-impermeable signalling molecules through α-hemolysin [7, 9, 14, 20, 21], or by synthesising a limited selection of membrane-permeable acyl homoserine lactones (AHSLs) used in bacterial quorum sensing (QS) [8, 10, 11]. QS systems are amongst the most simplistic modes of cellular communication. They involve as few as two proteins to function - one to produce the AHSL signalling molecule and another to detect it - making them ideal candidates for synthetic cell communication. While LuxI/R [8, 12], EsaI/R [10, 14], LasI/R [8], and RhlI/R [11] pathways have all been explored to date, they functioned at sub-optimal capacity since the substrates provided for AHSL biosynthesis were either homologues and less readily accepted by the AHSL synthases [11] or had to be assembled by accessory enzymes present in the cell lysate [8, 10]. For synthetic cells to become a mature technology with far-reaching applications in influencing the biology of living systems or as controllable delivery devices, a more diverse tool kit is necessary. In this work, we explored new synthetic and biological parts to expand synthetic cell capabilities in two specific areas; gene expression control and chemical communication.

First, we describe an alternative means of regulating gene expression that does not rely on naturally occurring small molecule-responsive transcription factors. Instead, gene expression inside the synthetic cells was controlled using ultraviolet (UV) light by incorporating light-sensitive 2-nitrobenzyl groups into the DNA templates. While this has previously been achieved by installing 1-(4,5-dimethoxy-2-nitrophenyl) diazoethane (DMNPE) groups into the backbone of plasmid-based DNA templates [22], we specifically installed the light-sensitive groups within the T7 promoters of linear DNA templates [23, 24]. The modified promoter had the same nucleobase sequence as the standard T7 promoter, except seven thymines were replaced by amino-C6-thymine bases to install photocleavable biotinylated (PCB) linkers into the major groove of the promoter region. Subsequent binding of monovalent streptavidin (mSA) to each biotin group created a steric blockade that impeded T7 RNA polymerase from binding to the T7 promoter. However, when exposed to UV light, the 2-nitrobenzyl group was photocleaved, and mSA liberated. Thus, *in situ* gene expression inside the synthetic cells was tightly repressed in the absence of UV light but restored after exposure to UV light. Here, LA-DNA templates were used to control *in situ* gene expression and remotely activate synthetic cells in a spatiotemporally-controlled manner.

Subsequently, we applied LA-DNA templates to control intercellular communication between synthetic and living cells. To this end, we reconstituted the BjaI/BjaR QS system from *Bradyrhozibium japonicum*, separately, in synthetic cells and *E. coli*. In this system, BjaI, an acyl CoenzymeA-dependent AHSL synthetase, was expressed inside the synthetic cells and synthesised a membrane-permeable signalling molecule, N-isovaleryl-L-homoserine lactone (IV-HSL), from its native membrane-impermeable substrates, isovaleryl CoenzymeA (IV-CoA) and S-adenosylmethionine (SAM). *E. coli* cells that expressed BjaR, an IV-HSL responsive transcription factor, acted as receiver cells in this communication network and coupled IV-HSL binding to the expression of a reporter gene held downstream of a BjaR-regulated promoter. We harnessed directed evolution of the BjaR transcription factor to confer improvements to the receiver cells and established a reporter system with a tight off state, high dynamic range (135-fold activation of gene expression), and exquisite sensitivity to IV-HSL (EC_50_=0.6 nM). This system is well suited for *in vitro* communication as the substrates for AHSL biosynthesis are commercially available and do not require additional enzymatic processing, the cognate AHSL contains a branched acyl chain making it structurally distinct from most other AHSLs, and gene expression is observed with as little as 100 pM IV-HSL [25].

These developments offer new ways to control synthetic cell activities and provide foundations for their application as targeted drug delivery devices and in studying communication in living systems.

## Results

### Light-activated gene expression inside synthetic cells

Genetically-controlled synthetic cells were constructed by placing the cellular machinery necessary for CFPS inside giant unilamellar vesicles (GUVs). This was accomplished by encapsulating an inner solution containing PURExpress, 200 mM sucrose, and Texas-Red dextran into egg phosphatidyl choline (Egg-PC) derived GUVs by emulsion droplet phase transfer [26]. First, the inner solution was emulsified in a lipid-containing oil to form water-in-oil emulsion droplets stabilised by a lipid monolayer. These emulsion droplets were then centrifuged through a second lipid monolayer formed at the interface between a lipid-containing oil and an outer solution (50 mM HEPES pH 7.6, 400 mM potassium glutamate, 200 mM glucose) to generate GUVs bound by a lipid bilayer (Figure S1). Gene expression within the synthetic cells was programmed according to linear DNA templates held within the inner solution, which would ultimately be modified to create the LA-DNA. These comprised a T7 promoter, ribosome binding site (RBS), gene of interest (GOI), and T7 terminator.

In the first instance, linear DNA templates encoding mNeonGreen (mNG) were selected, to visualise *in situ* gene expression using fluorescence microscopy. To enhance the cell-free expression of mNG inside the compartments, a T7 g10 leader sequence [27, 28] was introduced into the linear DNA templates. This sequence encoded the native 5’ untranslated region (UTR) and first 9 amino acids of the T7 bacteriophage gene 10 major capsid protein. In bulk CFPS reactions, DNA templates with the T7 g10 leader sequence expressed 1.6-fold more mNG than templates without the leader sequence (Figure S2a). Interestingly, inclusion of the N-terminal 9 amino acid coding leader alone, without the native 5’ UTR sequence, produced similar increases in mNG expression (Figure S2a). To elucidate whether these improvements were mNG specific, the T7 g10 leader sequence was also installed into linear DNA templates encoding mVenus. In this case, the leader sequence increased mVenus expression ~7-fold (Figure S2b). Thus, the placement of this N-terminal leader sequence upstream of a GOI may be a generalisable strategy for improving its expression in cell-free systems. Next, the linear mNG DNA containing the T7 g10 leader was introduced into synthetic cells. Cell-free expression of mNG proceeded rapidly within the first 2 hours of incubation at 37 °C and then continued to increase, albeit more slowly, for the proceeding 6 hours (Figure S3a). mNG expression within synthetic cells was somewhat heterogeneous, though this was expected due to the stochasticity of solute partitioning during emulsion phase transfer and the polydispersity of the GUVs formed [29]. In the absence of a DNA template mNG expression was not observed, confirming the synthetic cells were programmed according to the DNA template (Figure S3). In order to generate a richer, population level understanding of the synthetic cells, the diameters and mNG expression within the individual vesicles was quantified using a Matlab script (Figure S4; See supplementary data for code). mNG expression in individual 3-20 *µ*m synthetic cells was highly variable, spanning an order of magnitude at 8 hours (Figure S4b), and also tended to increase with GUV diameter (Figure S4c+d), as reported previously [30].

To control gene expression with light, photo-actuated T7 promoters were installed into the linear DNA templates by using a PCB-modified oligonucleotide as a primer in PCR [23]. Before LA-mNG DNA was introduced inside GUVs, we validated its assembly and performance using agarose gel electrophoresis and bulk CFPS reactions. DNA with PCB groups installed in the T7 promoter successfully bound mSA, forming LA-DNA. mSA remained stably fixed at the promoter until the 2-nitrobenzyl group within the PCB linkers underwent photolysis in a UV dose-dependent manner (Figure S5A). As expected, this tightly bound promoter-proximal blockade translated into a tight off-state when LA-mNG DNA was used for CFPS. When the reactions were not irradiated with UV light, minimal mNG expression was observed even after 4 hours incubation at 37 °C; gene expression was repressed by >97% relative to amine DNA templated reactions representing 100% photocleavage. In contrast, when the bulk CFPS reactions were exposed to UV light, mNG expression was recovered to ~66% of the amine DNA yield (Figure S5B).

Light-activated synthetic cells were assembled by encapsulating LA-mNG DNA inside the gene-expressing GUVs (Figure 1a). Minimal mNG expression was observed in the absence of UV light, and the fluorescence intensity of LA-DNA containing synthetic cells (-UV) was comparable to synthetic cells prepared without a DNA template. In contrast, when synthetic cells were irradiated with UV light, *in situ* mNG expression was recovered in >91% of synthetic cells. Similar to light-activated mNG expression in bulk, gene expression in the of LA-mNG synthetic cells (+UV) was restored to ~80% of that observed in synthetic cells containing amine mNG DNA (Figure 1b-c). Both the proportion of mNG-expressing vesicles and the fluorescence intensity of those vesicles increased in line with LA-DNA concentration (Figure S6), and as the UV light exposure time was increased from 5 to 10 minutes. However, beyond 10 minutes irradiation, mNG expression began to decrease (Figure S7). Thus, the LA-DNA templates retained sufficient blocking groups to prevent transcription at UV doses shorter than 10 minutes in duration, but gene expression was lower with prolonged UV light exposure as DNA/RNA/protein damage to the CFPS system exceeded any improvements afforded by the increased accessibility of T7 promoter sites.

**Figure 1:**
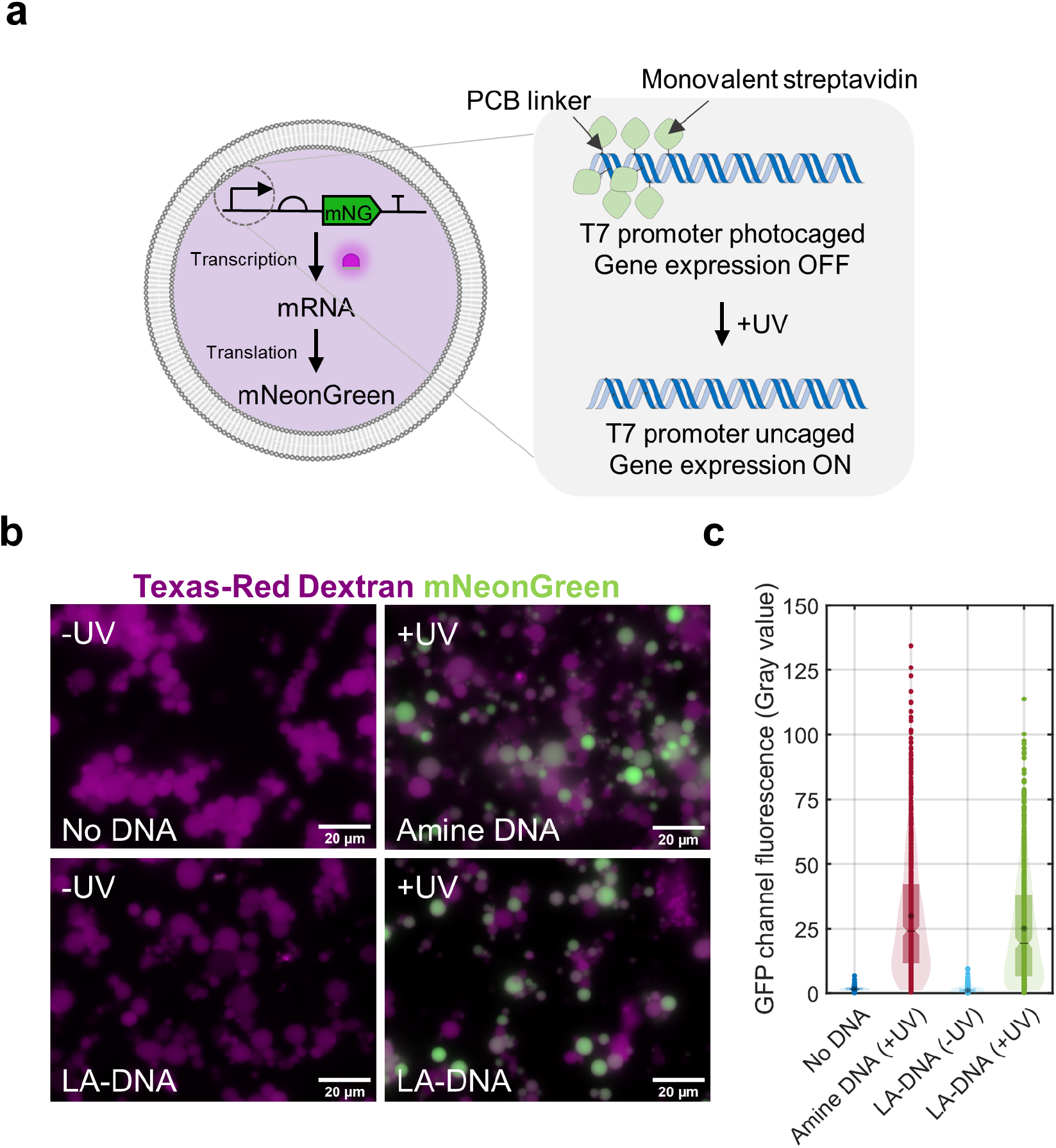
Light-activated gene expression in synthetic cells. **a)** PURExpress and LA-DNA templates were housed inside GUVs. In the absence of light, the photosensitive blockade prevented T7 RNA polymerase from binding the T7 promoter and transcribing the downstream gene of interest. After irradiation with UV light, T7 RNA polymerase could bind the uncaged T7 promoter to initiate the transcription, and subsequently translation of the protein of interest. **b)** Epifluorescence microscopy images of light-activated mNG expression inside synthetic cells with and without UV light exposure. Synthetic cells that were not irradiated with UV light expressed minimal mNG and their GFP channel fluorescence intensity was consistent with synthetic cells that contained PURExpress but no DNA template. UV-exposed synthetic cells expressed mNG, as demonstrated by the increase in green channel fluorescence. mNG fluorescence intensity was consistent with synthetic cells prepared with amine-modified mNG DNA templates. Scale bar = 20 *µ*m. **c)** Quantification of mNG expression in the individual GUVs using a circle detection based image analysis script. Fluorescence intensity was comparable across LA-mNG DNA (-UV) and no DNA samples, and the LA-mNG DNA (+UV) and amine mNG DNA (+UV) samples. (Median fluorescence intensity (No DNA) = 1.54 gray units (n=432); Median fluorescence intensity (LA-DNA (-UV)) = 1.09 gray units (n=568); Median fluorescence intensity (Amine DNA) = 25.2 gray units (n=627); Median fluorescence intensity (LA-DNA (+UV)) = 20.2 gray units (n=524)).

The major advantages of using light over small molecules to activate gene expression are that while the dosage is defined by the user in either case, light acts completely orthogonally to all small molecule-based regulation and can be applied in a spatially controlled manner. We exploited these traits to selectively activate gene expression inside light-activated synthetic cells. Patterned light used to activate gene expression was generated by projecting a collimated LED through a film photomask, onto light-activated synthetic cells held within imaging chambers (Figure 2a). The photomasks were designed to explore a range of simple and complex patterns, and contained features of varying dimensions to test the resolution limit of synthetic cell photolithography. LA-mNG DNA containing synthetic cells were held below the patterned light source inside a 1.5% ultra-low gelling agarose hydrogel. Immobilisation was necessary to prevent the GUVs from moving after light-activation and maintain the integrity of the patterned structures. Synthetic cells were irradiated with the patterned UV light for 10 minutes, and after 6 hours incubation at 37 °C, mNG was observed only in the synthetic cells residing within the light-exposed areas (Figure 2b). The fluorescent patterns generated by the mNG expressing synthetic cells closely matched the photomasks used to pattern the UV light. Simple photomask designs containing bulky features >200 *µ*m in size produced the best patterned structures, although, finer details down to ~100 *µ*m were also resolved in the more complex pattern designs (Figure S8). Overall, this confirmed gene expression inside GUVs could be remotely controlled with UV light, and specific populations of the light-activated synthetic cells could be activated via selective irradiation.

**Figure 2:**
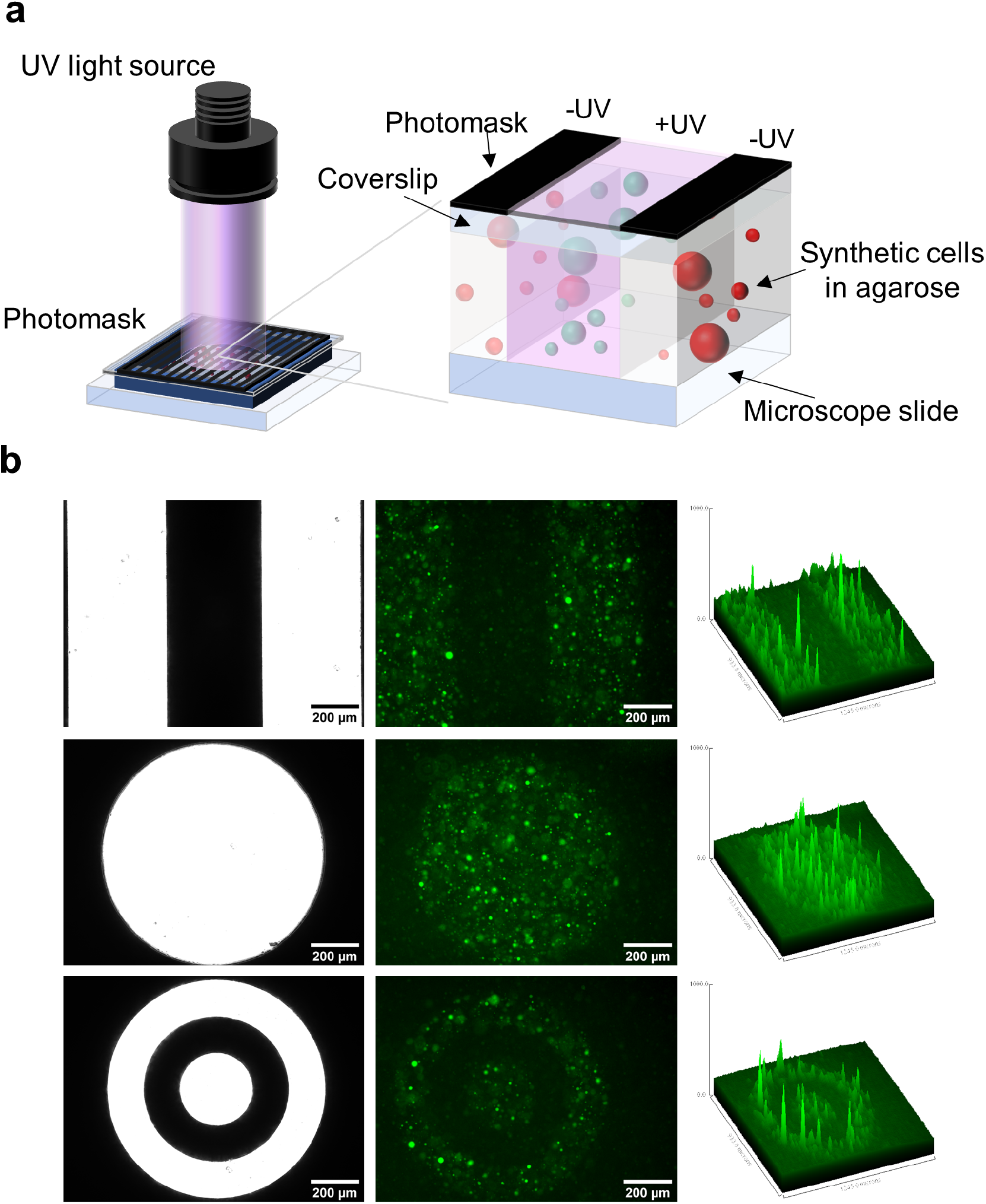
Spatially controlled activation of light-activated synthetic cells. **a)** Synthetic cells containing LA-mNG DNA were immobilised inside 1.5% agarose and gene expression was activated with photomask-patterned UV light. Dark regions of the photomask prevented the passage of light, and vesicles beneath did not express mNG. Lighter regions permitted the passage of light, uncaged the LA-DNA, and stimulated gene expression. **b)** Synthetic cells expressed mNG only in areas exposed to UV light. Patterns of mNG expression closely matched the photomask designs. Left: Photomasks, Centre: Epifluorescence microscopy images, Right: Surface plots. Scale bar = 200 *µ*m.

### Characterisation and optimisation of BjaR receiver cells

Next, we set out to establish a unidirectional communication pathway by reconstituting the BjaI and BjaR QS machinery, separately, within synthetic cells and *E. coli*, respectively. As a starting point, IV-HSL-responsive receiver cells were created by transforming *E. coli* XL10-Gold cells with a reporter plasmid encoding BjaR and GFP, pSB1A3 BjaR GFP [31]. BjaR was expressed from a constitutively active host promoter, while GFP expression was placed under the control of a pBjaR promoter formed by substituting the lux-box of the pLux promoter with the BjaR binding site (Figure S9). Hence, GFP should only be expressed when BjaR binds to IV-HSL and recruits an RNA polymerase to the pBjaR promoter (Figure 3a). Although this reporter plasmid had been characterised previously [31], GFP expression was initiated with N-butyryl-L-homoserine lactone (C4-HSL) rather than the cognate ligand, IV-HSL. Thus, we first evaluated the dose-response relationship between GFP expression and synthetic IV-HSL (synthesis described in supporting information). BjaR receiver cell cultures were grown in M9 minimal media to OD_600_ = 0.05, then transferred to a 96-well plate containing IV-HSL and incubated at 37 °C, 800 rpm, for 4 hours to initiate GFP expression. BjaR receiver cells expressed GFP in the presence of as little as 100 pM IV-HSL and expression saturated at ~100 nM IV-HSL, confirming the gene circuit was highly sensitive to IV-HSL (EC_50_ = 1.00 ± 0.11 nM). However, elevated GFP expression was observed in the absence of IV-HSL, and consequently, the reporter circuit had a poor dynamic range (3.6-fold activation) (Figure 3b). To elucidate the root cause of the high background fluorescence, a BjaR_KO_ variant of the reporter plasmid was constructed. Knocking out BjaR completely ablated GFP expression, both in the absence and presence of IV-HSL (Figure 3b). Therefore, the high OFF-state was attributed to BjaR-mediated transcription activation in the absence of IV-HSL, rather than host transcription factors recognising the pBjaR promoter, or transcription factor independent GFP expression.

**Figure 3:**
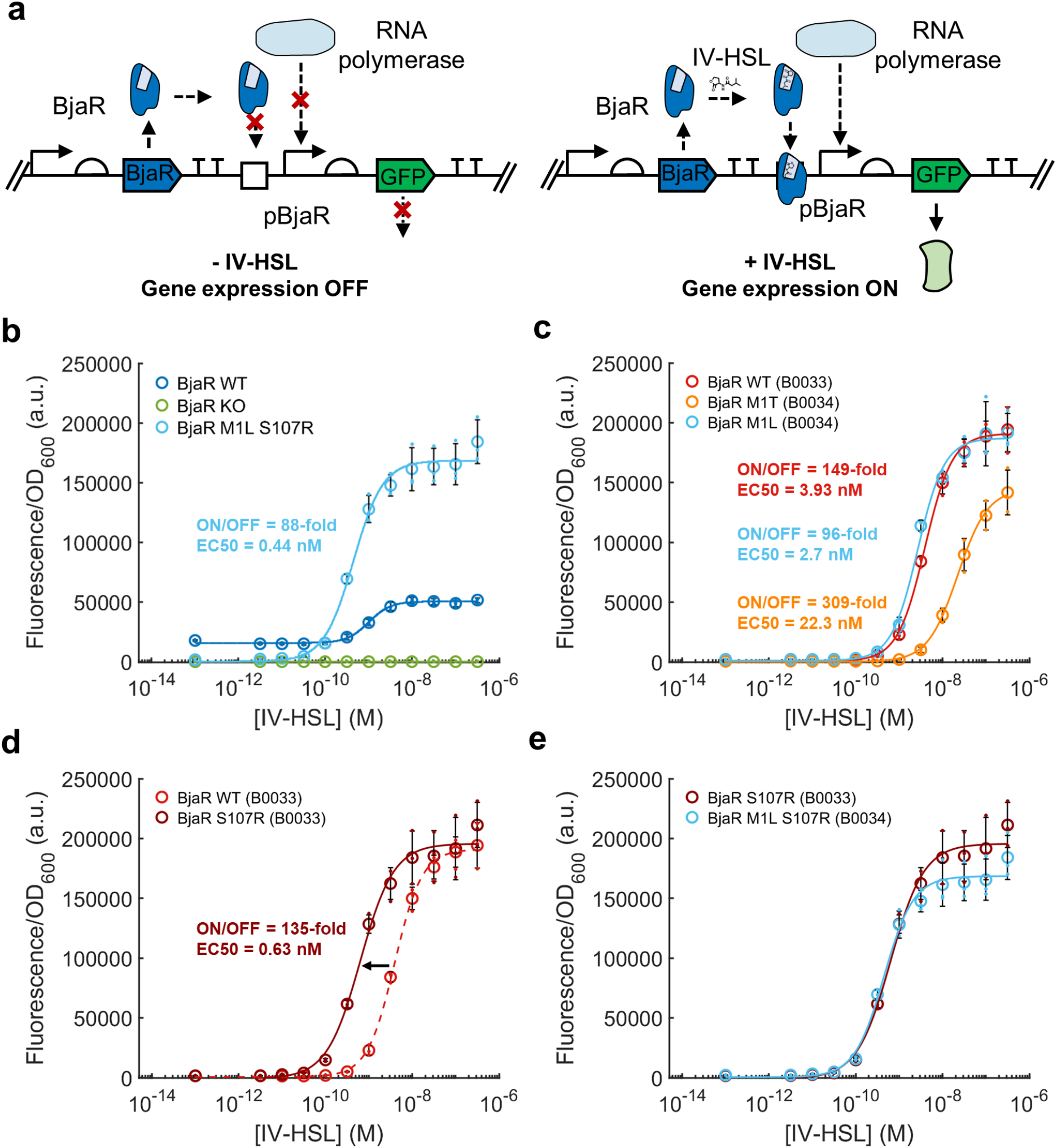
BjaR gene circuit is highly sensitive to IV-HSL. **a)** In the absence of IV-HSL, BjaR cannot recognise its consensus binding sequence, and GFP expression is turned off. On binding to IV-HSL, BjaR binds the operator site upstream of the pBjaR promoter, recruiting RNA polymerase and activating GFP expression. **b)** IV-HSL increased BjaR_WT_ receiver cell GFP fluorescence 3.4-fold, however, even in the absence of IV-HSL the receiver cells still expressed considerable amounts of GFP. BjaR_KO_ receiver cells showed baseline GFP expression in the absence of presence of IV-HSL. Cells containing the BjaR_M1L S107R_ mutant had a tighter off-state (lower baseline GFP expression) and an enhanced on-state (GFP expression in the presence of IV-HSL), as well as a marginal improvements in sensitivity to IV-HSL. **c)** The influence of the M1L mutation was reproduced by placing a weak RBS sequence upstream of the BjaR_WT_ gene sequence. An even tighter off state was observed for the M1T mutant, which affords a lower translation initiation rate. **d)** Receiver cells containing the S107R mutation were more sensitive to IV-HSL. **e)** Combining the B0033 RBS and S107R mutation in the BjaR gene closely replicated BjaRM1L S107R dose-response behaviour.

In pursuit of a BjaR reporter circuit with lower baseline GFP expression, we attempted to re-engineer the BjaR gene circuit towards stricter transcriptional control at the pBjaR promoter. We opted to use directed evolution with ON-state selection to generate BjaR mutants with a tighter OFF-state activity, as this had previously been used to improve the stringency of Lux-family transcription factors [32] (Figure S10a). Random mutations were introduced throughout the entire length of the BjaR gene by error-prone PCR; 100 *µ*M MnCl_2_ was added to Taq polymerase PCRs to introduce approximately two mutations into the gene per cycle [32]. The mutant BjaR gene library was then cloned into a modified reporter plasmid backbone containing both GFP (to score mutants) and the Kanamycin resistance gene (KanR) (for ON-state selection) downstream of the pBjaR promoter (Figure S10b). Transformants containing the BjaR mutant plasmids were grown in media supplemented with 100 *µ*g/mL ampicillin and 0.4-1 nM IV-HSL for 1 hour, to stimulate KanR (and GFP) expression, then for a further 5 hours in the presence of 200 *µ*g/mL kanamycin as a selection pressure. Cells that expressed functional BjaR variants, in turn, expressed KanR and survived ON-selection, whereas BjaR variants carrying deleterious mutations within the BjaR gene could not express KanR and were removed from the library. After ON-selection, the cells were washed and grown in media containing only ampicillin to enrich the population with fast-growing cells, i.e. those with lower cellular burden accomplished by more strictly controlling GFP and KanR expression. Plasmids from all cells present after outgrowth were then used as the DNA template in the next round of evolution. In total, BjaR underwent four rounds of random mutagenesis, ON-selection, and outgrowth. While directing the BjaR mutants towards strict expression control, we also ensured these new variants retained their intrinsically high sensitivity to IV-HSL by reducing the concentration of IV-HSL added to the cells by 0.2 nM for each subsequent round of evolution.

After four rounds of evolution, the ON and OFF states of 45 BjaR variants were screened by measuring their GFP expression in the absence and presence of 1 nM IV-HSL. 40 of these mutants demonstrated an improved dynamic range over the BjaR_WT_ protein, and 14 showed >25-fold activation of GFP expression (Figure S10c). Surprisingly, DNA sequencing of the top three performing mutants, colony 11, colony 28, and colony 36, revealed that they all contained the same mutations at Met1 (ATG-> CTG, Met->Leu) and Ser107 (AGC-> CGC, Ser-> Arg). In order to understand the true impact of these mutations on receiver cell performance, BjaR_M1L S107R_ was cloned back into the reporter plasmid backbone without KanR. Under this context, BjaR_M1L S107R_ demonstrated a ~88-fold activation of GFP expression granted by a 7.2-fold decrease in the OFF-state, and a 3.6-fold increase in the ON-state, and were also more sensitive to IV-HSL than BjaR_WT_ (EC_50_ BjaR_M1L S107R_ = 0.44 ± 0.04 nM) (Figure 3b).

Mutations at Met1 were unlikely to introduce novel functionality into the mutant protein. Rather, we reasoned that changes at the start codon would perturb the rate of translation initiation since other Met1 mutants with tight off-states were also identified in preliminary experiments [33]. Using a small library of ribosome binding sites (RBSs) with decreasing strengths to decouple the translation initiation rate from the identity of the initiation codon, we demonstrated that simply reducing the pool of BjaR available within the cells created receiver cells that mirrored the dose-response behaviour of the BjaR_M1L_ variant (Figure 3c & S11a). In fact, dose-response curves for BjaR receivers using the weakest RBS sequence tested (Bba B0033) and the M1L start codon were almost identical (Figure 3c) which is consistent with the notion that translation from the Bba B0033 RBS provides 0.1% of protein yield compared to the Bba B0034 RBS, just as an ACG start codon provides 0.1% of the protein yield compared to an ATG start codon [33, 34]. To elucidate the impact of the S107R mutation on gene circuit performance, we next introduced the S107R mutation alone back into the BjaR_WT_ gene, in the context of the Bba B0033 RBS background. The BjaR_WT_ (B0033) and BjaR_S107R_ (B0033) receiver strains had almost identical dose-response behaviours, except the BjaR_S107R_ mutant was ~ 6x more sensitive to IV-HSL than the BjaR_WT_ (EC_50_ BjaR_WT_ (B0033) = 3.93 ± 0.38 nM; EC_50_ BjaR_S107R_ (B0033) = 0.63 ± 0.10 nM) (Figure 3d). This is consistent with the location of Ser107 on the IV-HSL-binding domain [35]. Cells transformed with the BjaR_S107R_ (B0033) reporter plasmid granted the best combination of performance (135-fold activation range) and sensitivity (Figure 3e), and exhibited a 1.15-fold increase in the ON-state, plus a 1.33-fold decrease in the OFF-state compared to the BjaR_M1L S107R_ mutants obtained by directed evolution.

### Light-activated communication between synthetic cells and living cells

With an IV-HSL sensitive BjaR receiver cell in place, we turned our attention to creating IV-HSL producing synthetic cells. This was accomplished by expressing the BjaI AHSL synthetase within synthetic cells containing its native substrates, IV-CoA and SAM. IV-CoA (851.65 g/mol) and SAM (399.44 g/mol) are bulky, charged, membrane-impermeable molecules (logP_IV-CoA_ = −4.3; logP_SAM_ = −2.8), while IV-HSL produced by BjaI is smaller (185.22 g/mol) and more hydrophobic (logP_IV-HSL_ = 0.9) so passes freely across lipid bilayers. Hence, by expressing BjaI in GUVs containing SAM and IV-CoA, we anticipated IV-HSL would be synthesised *in situ* and then diffuse out of the synthetic cell, as is the case with other AHSLs generated in synthetic cells [8, 10, 11] (Figure 4A).

**Figure 4:**
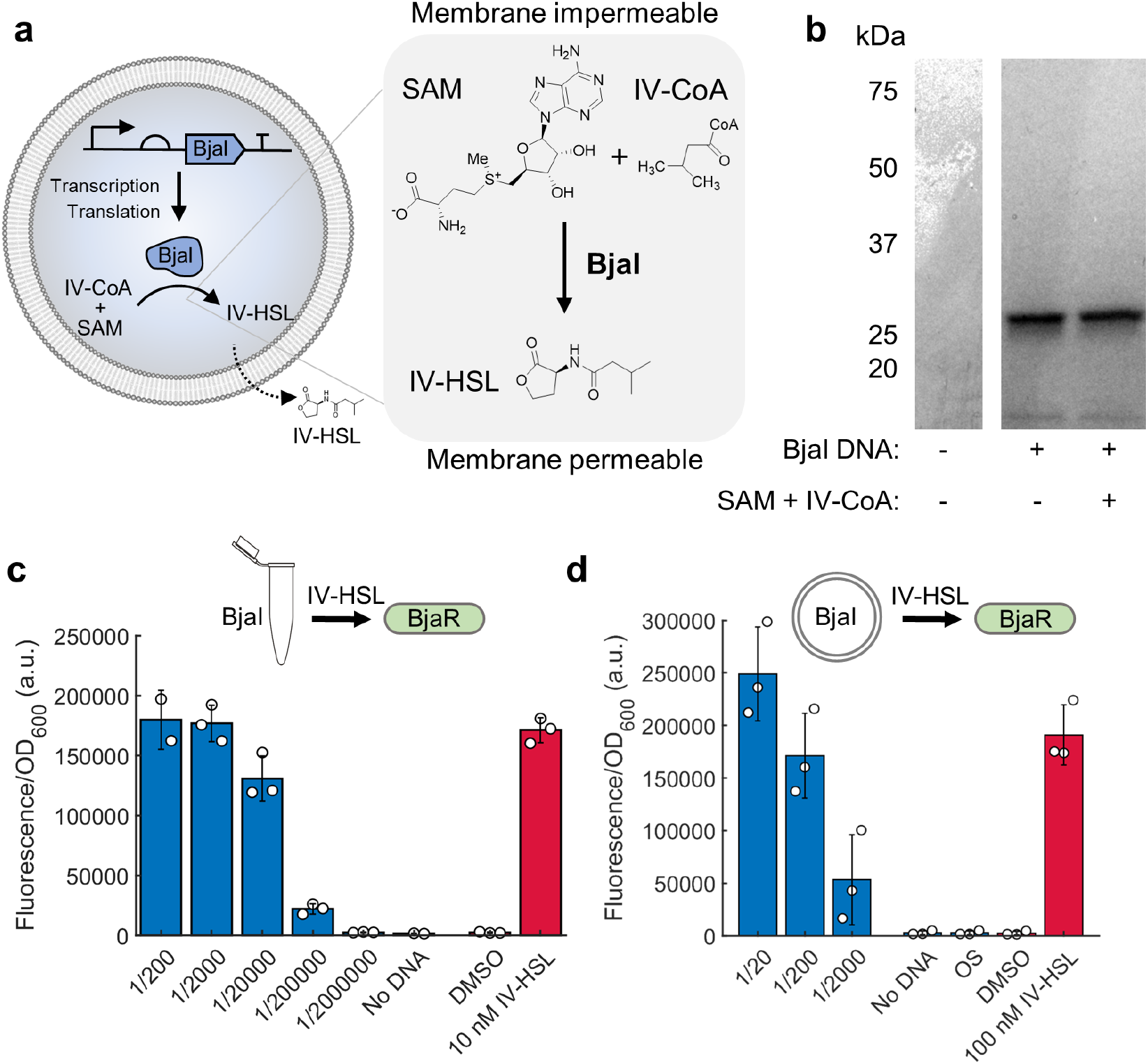
In situ IV-HSL biosynthesis by BjaI-expressing synthetic cells. **a)** IV-HSL biosynthesis by synthetic cells catalysed by the *in situ* expression of BjaI AHSL synthetase. Membrane impermeable substrates, IV-CoA and SAM, are converted into the membrane permeable signalling molecule IV-HSL. IV-HSL subsequently diffuses across the lipid bilayer into the external solution. **b)** Full length BjaI was expressed in bulk PURExpress reactions in the absence of prescence of IV-CoA and SAM. **c-d)** BjaI expressed in bulk CFPS reactions **(c)** and inside GUVs **(d)** was enzymatically active and synthesized IV-HSL from SAM and IV-CoA. 17.9 ± 2.6 *µ*M IV-HSL was produced by bulk BjaI IVTT reactions. 458.5 ± 119.6 nM IV-HSL was produced in synthetic cells. IV-HSL produced in bulk or by synthetic cells was quantified using BjaR _M1L S107R_ and BjaR _S107R_ (B0033) receiver cells, respectively.

We first validated BjaI expression using bulk CFPS reactions containing [^35^S]-methionine to visualise the newly synthesised protein. The full length BjaI protein (25 kDa) was successfully translated, and its expression was not influenced by the presence of SAM and IV-CoA (Figure 4b). To establish whether the expressed BjaI was enzymatically active, CFPS reactions containing 80 *µ*M IV-CoA and 300 *µ*M SAM were incubated at 37 °C for 5 hours, then serially diluted and incubated with BjaR receiver cells. Indeed, IV-HSL was successfully synthesised within the CFPS reactions as the reaction mixture containing the complete set of components (BjaI DNA + SAM + IV-CoA) fully activated GFP expression in the BjaR receiver cells, but no GFP expression was observed without BjaI expression (reactions containing SAM + IV-CoA, but no DNA) (Figure 4c). By comparing the fluorescence of the receivers in response to the biosynthetic IV-HSL with the dose-response curves obtained using the synthetic IV-HSL (Figure 3e), we estimated the CFPS reaction had a concentration 17.9 ± 2.6 *µ*M IV-HSL, thus 17.9 ± 2.6 pmol/*µ*L PURExpress was generated in the bulk CFPS reactions (Figure 4c). Similarly, on transferring this system inside GUVs, BjaI-expressing synthetic cells also produced sufficient IV-HSL to fully activate GFP expression. IV-HSL was present at 458.5 ± 119.6 nM in the 25 *µ*L synthetic cells samples, thus 2.28 ± 0.60 pmol/*µ*L inner solution was generated by the GUVs. The loss in specific activity between CFPS performed in bulk compared to inside GUVs was attributed to the loss of inner solution during emulsion phase transfer and when the GUVs ruptured, less efficient CFPS in GUVs compared to in bulk, and the fact not all of the GUVs were capable of gene expression.

Until now, synthetic cells had been prepared using 1X PURExpress and 200 mM sucrose, and had an osmolarity of ~1200 mOsm - far exceeding that of M9 minimal media (~300 mOsm). Before we could co-culture the synthetic cells with the receiver cells we first reduced the osmolarity of the inner solution to minimise the influx of water, GUV swelling, and membrane rupture that would ensue when transferring them into M9 miminal media. PURExpress is reported to tolerate slight deviation from its working concentration (around 20%), and so we tested IV-HSL production by BjaI expressed with PURExpress operating below the 1X working concentration. This confirmed that IV-HSL yields were only slightly perturbed at 0.4X the working PURExpress concentration in bulk (Figure S12a), IV-HSL was still produced by GUVs containing with 0.5X PURExpress (Figure S12b). GUVs prepared with an inner solution comprising 0.5X PURExpress without Murine RNase inhibitor, and 50 mM sucrose produced 32.5 ± 13.6 nM IV-HSL (0.16 ± 0.068 pmol/*µ*L inner solution). These synthetic cells were then interfaced with living cells by growing BjaR receiver cells on BjaI synthetic cell-laden agarose pads prepared with M9 minimal media. In this way, IV-HSL synthesized inside the GUVs by BjaI would passively diffuse across the lipid bilayer and into the agarose pad to activate GFP expression in the receiver cells. Despite the slight mismatch in osmolarity between the osmotically adjusted inner solution and M9 media, BjaI synthetic cells were stable in the agarose pads for at least 16 hours at 37 °C (Figure S13a), though GFP expression in the BjaR receiver was measured after 6 hours at 37 °C. BjaR_M1L S107R_ receiver cells were equally fluorescent when grown on M9 agarose pads containing 10 nM IV-HSL or BjaI synthetic cell-laden agarose pads, hence, IV-HSL was successfully synthesized and released into the agarose pad at concentrations sufficient to fully activate GFP expression (Figure S13b). Using this platform, we subsequently demonstrated remotely-controlled synthetic cell to living cell communication by encoding the BjaI gene on LA-DNA. BjaR receiver cells did not express GFP in the absence of UV light and had a background fluorescence equivalent to the no DNA control. Meanwhile, GFP expression was activated after irradiation with UV light, though the cells were less fluorescent than with 10 nM IV-HSL or amine BjaI synthetic cells, since IV-HSL yields were below the concentration threshold for full reporter gene expression (Figure 5).

**Figure 5:**
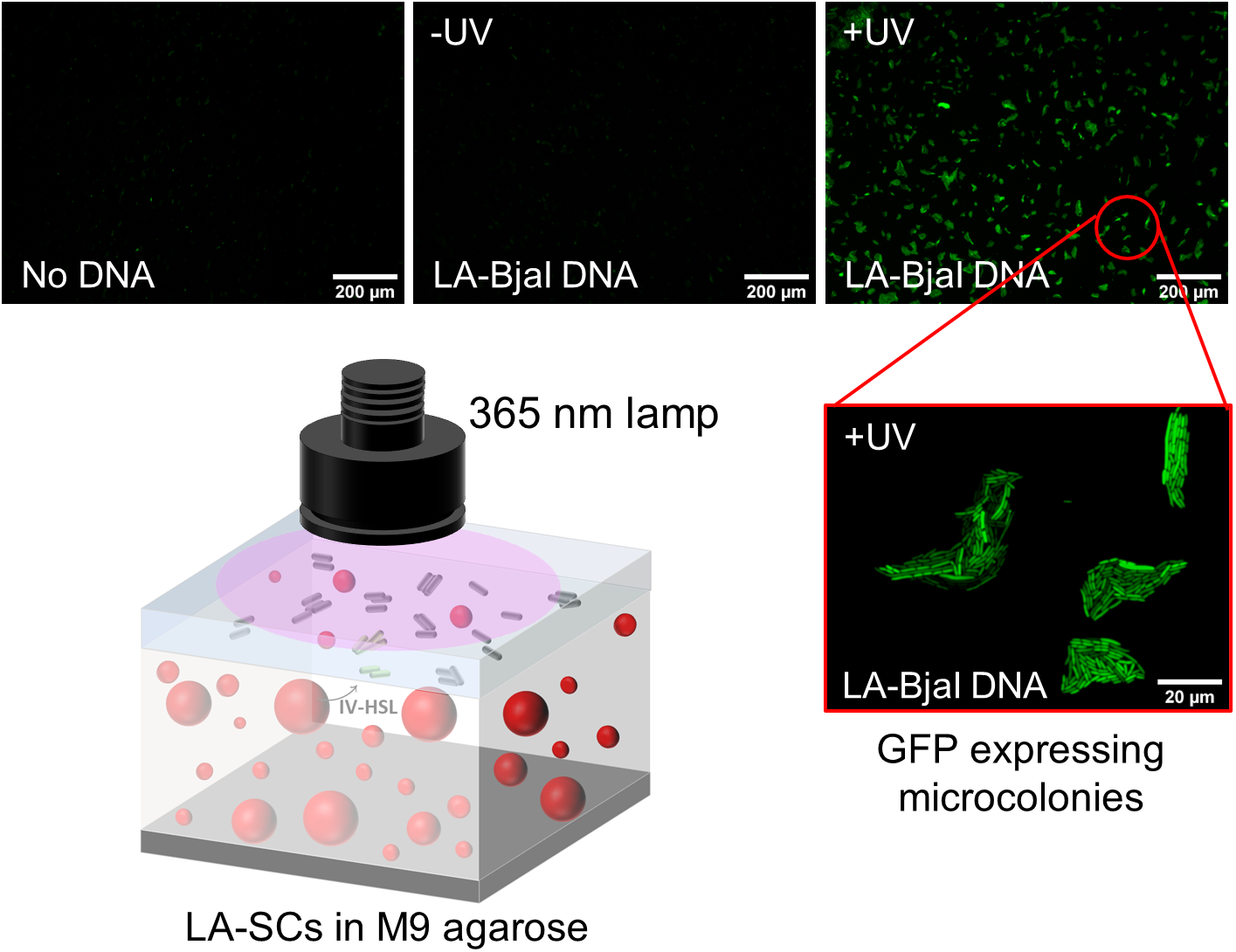
Light-activated communication between synthetic cells and living cells. IV-HSL-producing synthetic cells expressing BjaI were immobilised inside agarose pads containing M9 minimal media. IV-HSL-sensing receiver cells expressing BjaR were grown on top of the synthetic cell-laden agarose pads. After irradiation with UV light, IV-HSL produced inside the synthetic cells diffused into the agarose and activated GFP expression in the receiver cells held above. GFP expression was not observed in the absence of UV light, or without BjaI DNA templates present.

## Discussion

Synthetic cells are a versatile technology with the potential to serve as smart delivery devices or as a chassis for creating life from scratch. Despite the development of new tools and improvements in synthetic cell assembly methods, the biological parts used to regulate their activity have limited their reach to highly-controlled lab environments [15]. In the field’s preliminary work, well-established arabinose and IPTG-inducible transcription factors and theophylline responsive riboswitches were used to control *in situ* gene expression [7, 9]. Still, each performed poorly *in vitro* and represented a leaky, insensitive route of transcription/translation control that lacks physiological significance. Later, the transition to AHSL-sensitive transcription factors afforded synthetic cells the ability to sense and produce more biologically useful quorum sensing molecules, which are central to coordinating collective bacterial behaviours. While this marked considerable progress toward integrating synthetic cells with living cells, the most frequently adopted QS systems used to date, LuxR/LuxI and EsaR/EsaI, recognise and synthesise the same AHSL (3OC6-HSL), limiting the variety of synthetic cell activators that work orthogonally [8, 10, 12, 14].

In this work, we shifted away from the paradigm of using naturally derived parts to control gene expression. Instead, we utilised chemically modified LA-DNA templates to precisely control the location of synthetic cell activation with UV light. LA-DNA afforded a tight off-state in the absence of light, yet *in situ* gene expression was restored after exposure to UV light. This LA-DNA approach was implemented to regulate communication with *E. coli* cells using the BjaI/BjaR quorum sensing system; adding this unique branched AHSL into the synthetic cell communication toolbox. We believe this system is ideally suited to synthetic cell communication. It couples an acyl-CoA dependent synthetase, BjaI, which efficiently synthesises IV-HSL from its commercially available substrates, IV-CoA and SAM, with a highly sensitive IV-HSL-dependent transcription factor, BjaR, that activates gene expression at picomolar concentrations of IV-HSL. This strongly contrasts the properties of the RhlI/RhlR QS used previously in synthetic cells communication, which utilises a CoA substrate less readily than its native ACP-derived substrate [36], and requires >100 *µ*M C4-HSL to fully activate expression from pRhlI promoters [37]. To access the full capabilities of BjaR-regulated gene expression, we used directed evolution to identify mutations that improved the dynamic range of the receiver cells. Through this process, we identified mutations at the initiation methionine that drastically decreased the OFF-state and improved the ON-state, as well as a mutation at Serine107 (S107R) that made the receiver cells more sensitive to IV-HSL. While directed evolution is a powerful tool and explored the vast sequence space to uncover these mutations, an important lesson for the future is that this would not have been required if RBS strength was varied in the first instance; less can be more. That said, this finding validates the notion that there are multiple routes to achieving the same result, and suggests that simply decreasing RBS strength in other related AHSL-sensitive reporter plasmids [31] might enhance their performance. While this work was underway, another IV-HSL responsive gene expression was described that used a separate BjaR reporter plasmid system containing a mutated pBjaR promoter. It may be the case that this too reduced the potential of BjaR to activate the pBjaR promoter, through a version of the promoter that is recognised less readily [38]. By encoding the BjaI gene on the LA-DNA templates, IV-HSL biosynthesis was activated on exposing the synthetic cells to UV light. IV-HSL was membrane permeable and readily diffused across the lipid bilayer. We hoped to spatially pattern communication with living cells, however, this was not possible with the current set-up as expression occurred on a much longer timescale than molecular diffusion, and the membrane permeable nature of IV-HSL meant it was not retained within the cells.

While UV light was necessary to stimulate photocleavage of the 2-nitrobenzyl groups contained within the light-activated T7 promoters in this work, the modular nature of the LA-DNA templates makes them fully customisable. By installing longer wavelength photocages within the photocleavable biotin group, gene expression might be regulated according to more biocompatible and tissue penetrating visible or near-infrared light. In the future, light-activated synthetic cells might be used to study the mechanisms of cellular communication in a minimal environment or initiate communication pathways between living cells in a spatiotemporal manner. For instance, light-activated synthetic cells might be a useful tool for studying communication in biofilms by producing AHSLs or quorum quenching molecules originating at precise locations. Additionally, near-infrared-activated synthetic cells might be used as drug delivery devices that produce and secrete a therapeutic small molecule or protein at a target site.

## Materials and methods

### Materials

All consumables, solvents, and reagents were purchased from Sigma/Merck unless stated otherwise. Glass vials and teflon seal lids were purchased from Supelco. NHS-PCB-Biotin was purchased from Ambergen and stored in dry DMF at −80 °C. DreamTaq DNA polymerase mastermix, TexasRed-Dextran (10 kDa), and 25 *µ*L Gene frames were purchased from Thermofisher. PURExpress and all other enzymes were purchased from New England Biolabs (NEB). *E. coli* XL-10 Gold cells were purchased from Agilent. Egg PC was purchased from Avanti Polar Lipids. The 1 mm diamond drill bit was purchased from Eternal tools. The M365L2-C5 UV lamp was purchased from Thor Labs. Standard DNA oligonucleotides were synthesized by Integrated DNA Technologies. Photomasks were purchased from JD photodata. Amine-modified DNA oligonucleotides were synthesized by ATD-Bio. pMAT-mNeonGreen was synthesised by GeneArt. pSB1A3 BjaR-GFP was a gift from Prof. Karmella Haynes (Arizona State University). pET24a was a gift from Prof. Ben Davis (University of Oxford). Monovalent streptavidin was a gift from Prof Mark Howarth (University of Oxford). All DNA sequences can be found in Supplementary Table 1.

### Plasmids

mNG and BjaI genes were introduced into the PURExpress control template (now referred to as pPURE) in place of DHFR to create pPURE-mNG and pPURE-BjaI. pPURE-mVenus was prepared in previous work [23]. pPURE-T7g10-mNG and pPURE-T7g10-mVenus were prepared by replacing the 5’ UTR sequence of the pPURE plasmids with the T7 g10 leader sequence from [28], and introducing a 10 amino acid leader at the N-terminus of the respective genes. pPURE g10 mNG was prepared by introducing only the 10 amino acid leader sequence 5’ to the mNG gene. pSB1A3 BjaR_KO_ GFP was prepared by truncating the BjaR gene (BjaR_1-12_). pSB1A3 BjaR GFP KanR was prepared by placing the KanR gene, in frame, downstream of the GFP gene and pBjaR promoter. pSB1A3 BjaR_M1L S107R_ GFP and pSB1A3 BjaR_M1T_ were prepared by transferring the BjaR mutant genes from identified in the directed evolution screen back into the pSB1A3 BjaR GFP backbone. pSB1A3 (B0031-33) BjaR_WT_ GFP were prepared by replacing the B0034 RBS sequence with other RBS from the community library [34]. pSB1A3 BjaR_M1L_ GFP and pSB1A3 (B0033) BjaR_S107R_ GFP were prepared by site directed mutagenesis. Plasmid assembly is described, in full, in the supplementary details.

### Preparation of linear DNA templates

Linear DNA templates were obtained by PCR of pPURE plasmids encoding the desired gene of interest. PCRs were performed with Phusion DNA polymerase mastermix using 1 ng plasmid and 500 nM T7 for/rev1 primers. Reactions were cycled according to the following programme: 98 °C for 30s, 35 cycles of [98 °C for 10 s, 54 °C for 20 s, 72 °C for 30 s], 72 °C for 5 minutes, 4 °C HOLD. PCR products were validate by agarose gel electrophoresis and purified using a QIAquick PCR purification kits. DNA templates were ethanol precipitated and resuspended in Milli-Q H_2_O to 75-100 ng/*µ*L.

### Modification of amine DNA

The PCB-modified forward primer was prepared as described previously [23]. 10 *µ*M T7 For Amine was combined with PCB-NHS (Final conc = 5 mM), 100 mM sodium bicarbonate and DMF (Final conc = 50% v/v) to a final volume of 50 *µ*L in 500 *µ*L protein-lo bind tubes and incubated in the dark for 3 hours with gentle shaking. After 3 hours, reactions were quenched with 5 *µ*L 1 M Tris pH 8.0. 445 *µ*L Milli-Q H_2_O were added to the tube, and then transferred to a 0.5 mL 3 kDa Amicon column. Samples were centrifuged at 14,000 x g for 15 mins, topped up to 500 *µ*L with 10 mM Tris pH 8.0, then centrifuged again (3 times in total). 2 × 50 *µ*L samples were injected onto a Discovery C-18 column pre-equilibrated with 5% ACN, 95% buffer A (10 mM ammonium acetate) and separated with a 5-35% buffer A:ACN gradient over 40 mins, flow rate 2 mL/min. Fractions corresponding with the major peak were pooled, lyophilised, redissolved in 500 *µ*L Milli-Q H_2_O and lyophilised again to remove trace ammonium acetate. Lyophilised oligos were made up in 500 *µ*L Milli-Q H_2_O, and washed 4 times in a 0.5 mL 3 kDa Amicon centrifugal filter column.

### Formation of light-activated DNA

PCB-DNA templates were prepared by PCR with DreamTaq DNA polymerase master mix, 325 nM T7 For PCB/250 nM Rev1 primers, and 1 ng DNA template with the following thermal cycler programme: 95 °C for 2 mins, 35 cycles of [95 °C for 30 s, 49 °C for 30 s, 72 °C for 1 min], 72 °C for 5 mins, 4 °C HOLD. PCR products were purified using the QIAquick PCR purification kit, ethanol precipitated and resuspended in Milli-Q H_2_O. 1 *µ*g of linear PCB-DNA was added to a 50x molar excess of monovalent streptavidin and made up to 50 ng/*µ*L with 10 mM tris pH 8.0. DNA was incubated with mSA at 24 °C, 600 rpm for 3 hours in the dark. Samples were then kept at 4 °C for at least 12 hours before use. Amine-modified DNA representing the 100% photocleaved LA-DNA was prepared in the same way, except the T7 for amine primer was used in place of T7 for PCB.

### Gel electrophoresis of LA-DNA

50 ng of LA-DNA was irradiated with a collimated 365 nm UV lamp (Thor Labs, M365L2-C5) for 0-15 minutes at 0.75 mW, then combined with 6X loading dye and loaded into wells of a 1.5% TAE agarose gel. Gels were run in 1X TAE buffer at 100 V for 1 hour 15 mins, then stained in 3X GelRed for up to 1 hour. Gels were imaged using a BioRad Geldoc XR+ gel imager.

### Bulk CFPS of fluorescent proteins

3 *µ*L IVTT reactions were prepared with 1X PURExpress and supplemented with 0.8 U/*µ*L Murine RNase inhibitor, and 5 ng/*µ*L linear mNG/mVenus DNA template or 7.5 ng/*µ*L amine/LAmNG DNA. Where indicated, reactions were irradiated with UV light (0.75 mW) for 1-5 mins. Reactions were incubated in a thermal cycler at 37 °C for 4 hours, then held at 4 °C for at least 15 minutes. Reactions were worked up with 47 *µ*L PBS and transferred to Perkin Elmer 384-well black flat bottom optiplates. Fluorescence was measured using a Tecan Infinite M1000 fluorescence plate reader. Bandwidth = 5 nm; Z-position = 23179 *µ*m; Ex_mNG_ = 506 nm; Em_mNG_ = 517 nm; Ex_mVenus_ = 515 nm; Em_mVenus_ = 528 nm.

### Imaging chambers

1 mm holes were drilled through 25 mm x 75 mm microscope slides using a 1 mm diamond toolbit and Dremel 3000 rotary tool to correspond with diagonal corners of 25 *µ*L gene frames. All slides and coverslips (22 mm x 22 mm) were cleaned with 2% decon, isopronaol and Milli-Q H_2_O, then sonicated in Milli-Q H_2_O for at least 10 minutes and dried under N_2_ flow before use. Coverslips were O_2_ plasma treated for 5 minutes, before 25 *µ*L gene frames were attached. 0.1% BSA in PBS was added inside the gene frames and incubated at room temperature for 15 minutes. BSA was removed and coverslips were washed with outer solution, twice. Outer solution was removed and gene frames were applied to the drilled microscope slides. 25 *µ*L of samples were introduced through the drilled holes and then the holes were sealed with double sided tape. Samples were incubated coverslip side down. See Figure S14 for illustration.

### Assembly of mNG and LA-mNG synthetic cells

2 mL glass vials were cleaned with isopropanol and then held under vacuum for > 1 hour. Egg PC dissolved in chloroform (50 mg/mL stock) was transferred to the vial using Hamilton syringes, then the held under a gentle stream of N_2_ to evaporate the chloroform. The vials were tilted at 45^*o*^ and rotated slowly while held under N_2_ flow to distribute the lipids evenly up the walls. Vials containing lipid films were held under vacuum in a desiccator for 2-3 hours to remove residual chloroform. Mineral oil (filtered through 0.22 *µ*m PES membrane) was added to Egg PC films, by mass, to a final concentration of 5 mg/mL Egg PC. Vials were vortexed for 1 minute, then incubated in a heat block at 80 °C for 10 minutes with the lids removed. Lids were reapplied to vials and sealed tightly using Teflon tape and parafilm, then vortexed aggressively for 1 minute and sonicated in a sonication bath heated to 50 °C for 1 hour. Lipid containing oil was stored at room temperature overnight and vortexed, then sonicated for 10 mins at room temperature immediately before use.

250 *µ*L of 5 mg/mL Egg PC in mineral oil was transferred to 1.5 mL centrifuge tubes and placed on ice. 5 *µ*L inner solution (PURExpress containing 5-10 ng/*µ*L DNA, 0.8 U/*µ*L Murine, 25 *µ*M TR-Dextran (10 kDa) and 200 mM sucrose) was added into the chilled lipid-containing oil ensuring the tip was constantly moved through the lipid-containing oil to disperse the inner solution. Tubes were passed across a centrifuge rack 3-5 times using light pressure to form cloudy W/O emulsions that were place on ice for ~5 minutes. Meanwhile, 100 *µ*L of lipid-containing-oil was layered on top of 250 *µ*L of chilled outer solution (50 mM HEPES, 400 mM potassium glutamate, 200 mM glucose (pH 7.6)) and place at room temperature. W/O emulsions were then added on top of this oil phase, and this tube was placed on ice for ~5 minutes. Centrifuge tubes containing the W/O emulsion above outer solution were centrifuged at 16,000 x g, 4 °C, for 30 minutes.

After centrifugation, the oil phase and ~200 *µ*L of outer solution was removed from the tube and discarded. Using a fresh tip, ~10 *µ*L of the remaining outer solution was ejected against the GUV pellet to displace it from the tube, and the intact GUV pellet was transferred to a new tube containing 250 *µ*L outer solution. The pellet was subsequently resuspended by gently pipetting up and down. Vesicles were centrifuged at 10,000 x g, 4 °C for 10 minutes, then the outer solution was removed and the pellets were gently resuspended in 25 *µ*L of fresh outer solution.

### mNG expression in synthetic cells

To assess mNG expression from inside the synthetic cells, the GUV pellets were resuspended in 25 *µ*L outer solution and loaded into home-made imaging chambers. Synthetic cells were incubated in a 37 °C oven for 2-8 hours, coverslip side down. Where indicated, LA-mNG SCs were irradiated with UV light for 0-12.5 mins at 0.75 mW. Fluorescence microscopy was performed using a a Lecia DMi8 inverted epifluorescence microscope using a 100x oil immersion objective lens. GUVs were imaged using the brightfield, TXR, and GFP filters.

### Patterning mNG expression in synthetic cells

The pelleted LA-mNG GUVs were resuspended in 15 *µ*L molten 1.5% ultra low gelling point agarose prepared with outer solution, maintained at 28 °C and added into a 25 *µ*L gene frame sandwiched between a glass coverslip and microscope slide. Samples were gelled by placing them at 4 °C for ~30 minutes. Film photomasks were placed on top of the coverslips and irradiated under the UV lamp for 10 minutes at 0.75 mW (Power adjusted to account for the transparent film containing photomask designs). Samples were incubated at 37 °C for 6 hours then imaged with a Lecia DMi8 inverted epifluorescence microscope using a 10x onjective lens. GUVs were imaged using the brightfield, TXR, and GFP filters.

### Image processing

TXR and GFP channel brightness was normalised across all images within a single experiment, then the separate channels were saved as individual .PNG files. All images corresponding to a single sample were stored within the same directory. ‘Background’ images were created by manually selecting vesicle-free regions of microscopy images (1 from each sample within the experiment) and measuring the mean pixel intensity. PNG files were input into the vesicle analysis script (see appendix for scripts). Manual analysis of GUVs was performed in ImageJ by creating a region of interest corresponding with the area of an individual vesicle using the circle tool and measuring the diameter and mean pixel intensity within this ROI for both the TXR channel and GFP channel images. Values were exported into excel file and plotted using matlab.

### IV-HSL synthesis

To a stirring suspension of L-Homoserine lactone hydrochloride (100 mg, 0.726 mmol) in acetonitrile (5 mL, 0.145 mmol) was added diisopropylethylamine (0.32 mL, 1.8 mmol, 2.5 eq.). Isovaleryl chloride (0.14 mL, 1.4 mmol, 1.9 eq.) was then added drop wise over 10 minutes and the reaction left stirring for 16 hours. The resulting brown solution was then concentrated *in vacuo* and subjected to column chromatography (1:1 to 0:1 PE_40-60_:EtOAc) to yield a white, crystalline solid as the desired product (78 mg, 0.422 mmol 58%). ^1^H-NMR (400 MHz, CDCl_3_) 5/ppm 5.99 (NH, br s, 1H), 4.54 (lac CH, ddd, *J* = 11.6, 8.6, 5.7 Hz, 1H), 4.47 (lac CH, td, *J* = 9.1, 1.3 Hz, 1H), 4.28 (lac CH, ddd, *J* = 11.2, 9.3, 5.9 Hz, 1H), 2.88 (lac CH, dddd, / = 12.5, 8.6, 5.8, 1.2 Hz, 1H), 2.21 - 2.02 (tail CH_2_ + tail CH + lac CH, m, 4H), 1.02 - 0.92 (CH_3_, m, 6H). MS (ESI^+^) found 208.2, [M+Na]^+^ requires 208.1. Data in accordance with literature [39].

### M9 minimal media

Supplemented M9 minimal media (1X M9 salts, 0.34 mg/mL thiamine hydrochloride, 0.2% Cas amino acids, 2 mM MgSO_4_, 100 *µ*M CaCl_2_, 0.4% (w/v) glucose) [40] was prepared fresh for every experiment. Media was sterilised by passing it through a 0.22 *µ*m PES syringe filter before use.

### Characterising BjaR receiver cells

BjaR receiver cells were incubated in 5 mL LB + Amp media overnight. Overnight cultures were diluted 1/250 or 1/500 in 5 mL M9 minimal media + Amp and incubated at 37 °C, 225 rpm for 3 hours 45 mins (until OD_600_ ~ 0.075). Cultures were diluted down to OD600 = 0.05 with M9 minimal media + Amp and 200 *µ*L was added to wells of a 96-well plate containing 1 *µ*L IV-HSL dissolved in DMSO or 1 *µ*L BjaI CFPS reactions (IV-HSL concentrations obtained from serial dilution from 63.2 *µ*M to 632 pM; final DMSO concentration = 0.5%). Cells were incubated at 37 °C, 800 rpm in a thermomixer with a heated lid for 4 hours. Fluorescence and OD_600_ were measured in a Tecan M1000 infinite plate reader (Ex: 488 nm, Em: 510 nm, BW: 5 nm Gain: 101, Z-position: 21311 *µ*m).

### Directed evolution

BjaR underwent 4 rounds of directed evolution based on a previously published protocol [32]. Briefly, epPCR of the BjaR gene was performed in the presence of 100 *µ*M MnCl_2_ using Taq DNA polymerase and 500 nM pSB1A3 for 1 and pSB1A3 rev 1 primers. Reactions were cycled using the following programme: 95 °C for 30 s, 30 cycles of [95 °C for 15 s, 51 °C for 30 s, 68 °C for 1 min], 68 °C for 5 mins, 4 °C HOLD. PCR products were electrophoresed on a 1.2% TAE agarose gel and gel extracted using a Monarch Gel extraction kit. Gel extracted BjaR DNA was cloned into the pSB1A3 GFP-KanR BB using a home-made Gibson’s assembly master mix [41] and transformed into *E. coli* XL10-Gold. Cells were incubated in 500 *µ*L SOC media for 1 hour at 37 °C, 225 rpm, then 4.5 mL LB + Amp (100 *µ*g/mL final) was added, and cells were incubated overnight at 37 °C, 225 rpm. 1 × 10^7^ cells from overnight culture were transferred into 1 mL M9 minimal + 100 *µ*g/mL Amp in the presence of IV-HSL (1 nM/0.8 nM/0.6 nM/0.4 nM for rounds 1-4 respectively) and incubated for 1 hour 37 °C. After 1 hour, kanamycin was added to a final concentration of 200 *µ*g/mL and incubated for a further 5 hours. Cells were centrifuged at 8,000 x g for 3 mins to remove the selective media, then resuspended in 1 mL LB, repelleted, and resuspended in 2 mL LB + Amp. Cells that survived ON-selection were incubated for 2 hours, 37 °C. Cells were pelleted and plasmids extracted using a PUREyield mini prep kit. Plasmids from rounds 1/2/3 were used as DNA templates for epPCR in rounds 2/3/4, respectively. After 4 rounds of evolution, 45 mutants were picked and used to inoculate 500 *µ*L LB + 100 *µ*g/mL Amp in 2 ml deep well plates. Cells were incubated at 37 °C, 250 rpm overnight, then 5 uL of cells were used to seed 500 *µ*L M9 minimal media + 100 *µ*g/mL Amp in the remaining wells of the deep well plate. Cells were grown at 37 °C for 3/4 hours, then 200 *µ*L were transferred to a lidded, transparent corning 96 well microplate containing either DMSO or 1 *µ*L 200 nM IV-HSL. Plates were incubated at 37 °C, 800 rpm in a thermomixer with a heated lid for 4 hours. Fluorescence and OD_600_ were measured in a Tecan M1000 infinite plate reader (Ex_GFP_: 488 nm; Em_GFP_: 510 nm; BW: 5 nm; Z-position: 21311 *µ*m).

### Bulk CFPS of BjaI [^35^S]-Methionine labelled

3 *µ*L CFPS reactions were prepared using PURExpress with 0.8 U/*µ*L Murine RNase inhibitor, 5 ng/*µ*L linear BjaI DNA template, and [^35^S]-Methionine (0.3 *µ*L, 1,200 Ci mmol^-1^, 15 mCi mL^-1^, MP Biomedicals). SAM and IV-CoA were added to 300 *µ*M + 80 *µ*M, respectively, where indicated. Reactions were incubated in a 37 °C water bath for 3 hours, then run on a precast 10% mini protean TGX gel at 200 V for 30 mins. The gel was dried onto filter papers using a Biorad Model583 gel dryer, exposed to Kodak Biomax MS film overnight and then developed.

### IV-HSL biosynthesis by BjaI

3 *µ*L CFPS reactions were prepared using PURExpress with 0.8 U/*µ*L Murine RNase inhibitor, 5 ng/*µ*L linear BjaI DNA template, and 300 *µ*M SAM and 80 *µ*M IV-CoA, where indicated. Reactions were incubated for 5 hours at 37 °C using a thermal cycler, then serially diluted in MilliQ-H_2_O. 1 *µ*L of the samples was added to lidded Corning 96-well plates containing 200 *µ*L BjaI receiver cells (pSB1A3 BjaR_M1L S107R_) at OD_600_ = 0.05. Cells were grown at 37 °C, 800 rpm in a thermomixer with a heated lid for 4 hours. OD_600_ and fluorescence were measured using a Tecan infinite M1000 plate reader. Ex = 488 nm; Em = 510 nm; Bandwidth = 5 nm; Z-position = 23179 *µ*m; Gain = 101.

### BjaI expression in synthetic cells

Synthetic cells were assembled as described above, except the inner solution was prepared without TexasRed-Dextran and contained 10 ng/*µ*L linear BjaI DNA, 80 *µ*M IV-CoA, and 300 *µ*M SAM. Samples were resuspended in 25 uL outer solution and incubated in centrifuge tubes at 37 °C, 250 rpm for 5 hours. The BjaI synthetic cells were then serially diluted and assayed against BjaI receiver cells, as described above. In synthetic cells prepared at 0.75X and 0.5X PURExpress concentrations and 50 mM sucrose, the outer solution was adjusted accordingly (50 mM HEPES pH 7.6, 300 mM potassium glutamate, 50 mM glucose and 50 mM HEPES pH 7.6, 200 mM potassium glutamate, 50 mM glucose, respectively).

### Agarose pad induction of receiver cells

Microscope slides were first cleaned with Milli-Q H_2_O and isopropanol. 2 × 25 *µ*L gene frames were attached to a single microscope slide with the open side still covered with the transparent film. Molten 1.5% ultra-low gelling point agarose was mixed with SCs or 10 nM IV-HSL as required and added inside the gene frames. A second microscope slides was carefully slid on top of the gene frames to distribute the molten agarose throughout the frame. Microscope slides sandwiching the gene frame were transferred to the fridge for 1 hour to gel the agarose, then the top slide was carefully removed by moving it off of the gene frame horizontally. 3 *µ*L BjaR_M1L S107R_ receiver cells at OD_600_ = 0.1 were added on top of the gelled agarose pads and the excess liquid was allowed to evaporate for ~5 mins. The transparent film was removed from the gene frame and an isopropanol cleaned, plasma treated coverslip was placed on top of the agarose pad to seal it. Samples were irradiated with 0.75 mW UV light for 5 minutes where indicated, then incubated at 37 °C for 6 hours and imaged using a with a Lecia DMi8 inverted epifluorescence microscope using a 10x onjective lens.

## Acknowledgements

We would like to thank Prof M. Howarth and Dr M. Fairhead for providing the monovalent streptavidin, Prof. K Haynes for providing the pSB1A3 BjaR_WT_ GFP plasmid, and Prof. Ben Davis for the pET24a plasmid.

J.M.S. is supported through the Synthetic Biology Centre for Doctoral Training - EPSRC funding (EP/L016494/1).

D.H. is grateful to the EPSRC Centre for Doctoral Training in Synthesis for Biology and Medicine (EP/L015838/1) for a studentship, generously supported by AstraZeneca, Diamond Light Source, Defence Science and Technology Laboratory, Evotec, GlaxoSmithKline, Janssen, Novartis, Pfizer, Syngenta, Takeda, UCB and Vertex.

M.J.B. is supported by a Royal Society University Research Fellowship and an EPSRC New Investigator Award (EP/V030434/1).

## Contributions

M.J.B and J.M.S designed the project. J.M.S designed, performed, and analysed the experiments. D.H synthesized the IV-HSL. J.M.S and M.J.B wrote the paper.

