## Supplementary Information for "Engineering cellular communication between light-activated synthetic cells and bacteria"

#### 8 Supplementary data

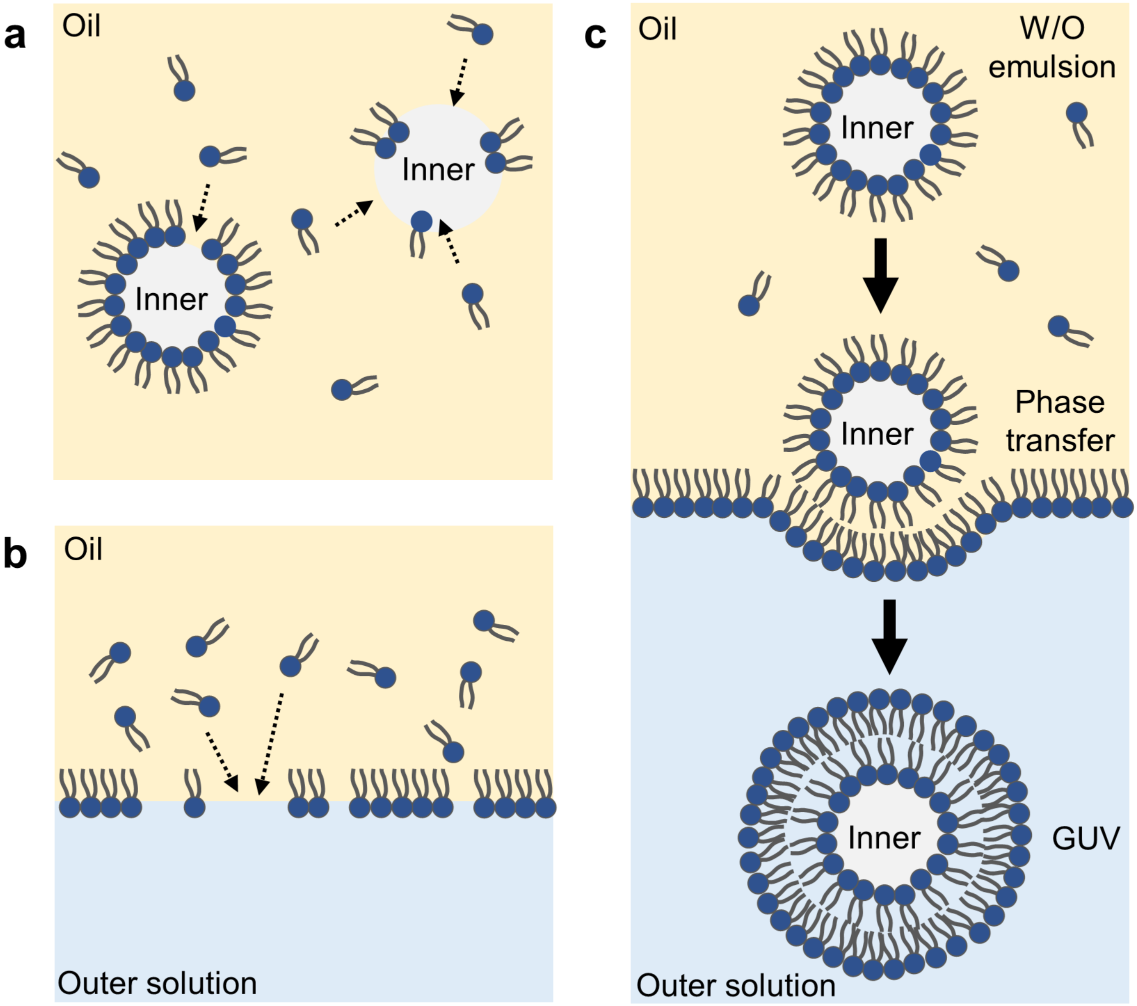


**Figure S1: Formation of GUVs by emulsion droplet transfer**
**a)** An aqueous sucrose-containing inner solution was emulsified in a lipid-containing oil to create creating droplets stabilised by a lipid monolayer. **b)** Lipid-containing oil was placed on top of an aqueous solution containing glucose to create a second lipid monolayer at the W/O interface. **c)** Emulsion droplets were pulled through the second lipid monolayer into the outer solution by centrifugation.

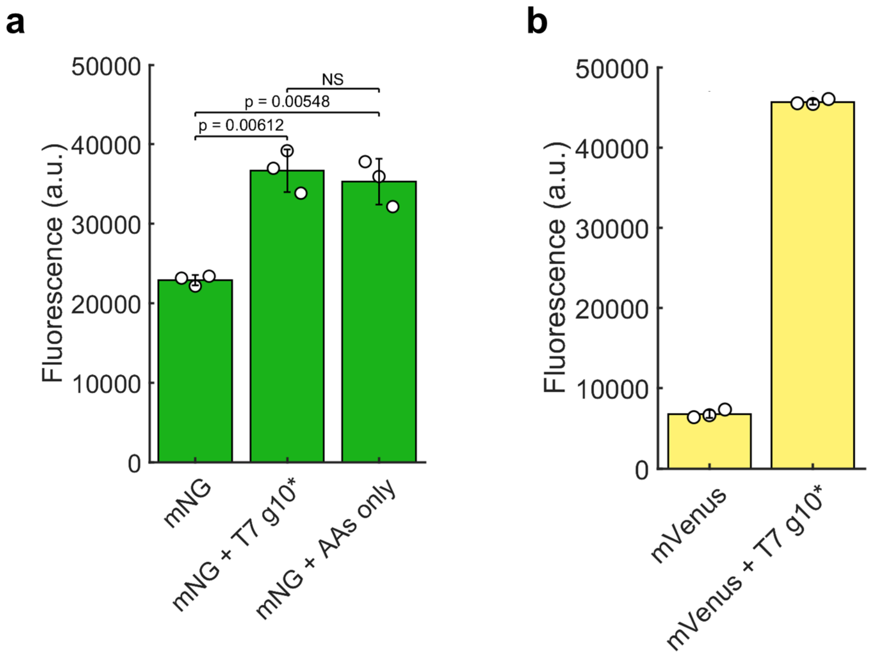


**Figure S2: Improving the CFPS of mNG**
**a)** Installing the T7 g10 leader into the mNG linear DNA template increased mNG fluorescence intensity 1.6-fold. The same improvements in mNG expression were also observed when only the 9 amino acid leader (MASMTGGQQ; ATGGCTAGCATGACTGGTGGACAGCAA) was introduced at the 5' end of the mNG gene. **b)** Inclusion of the T7 g10 leader sequence into linear DNA templates encoding mVenus improved its expression 7-fold.

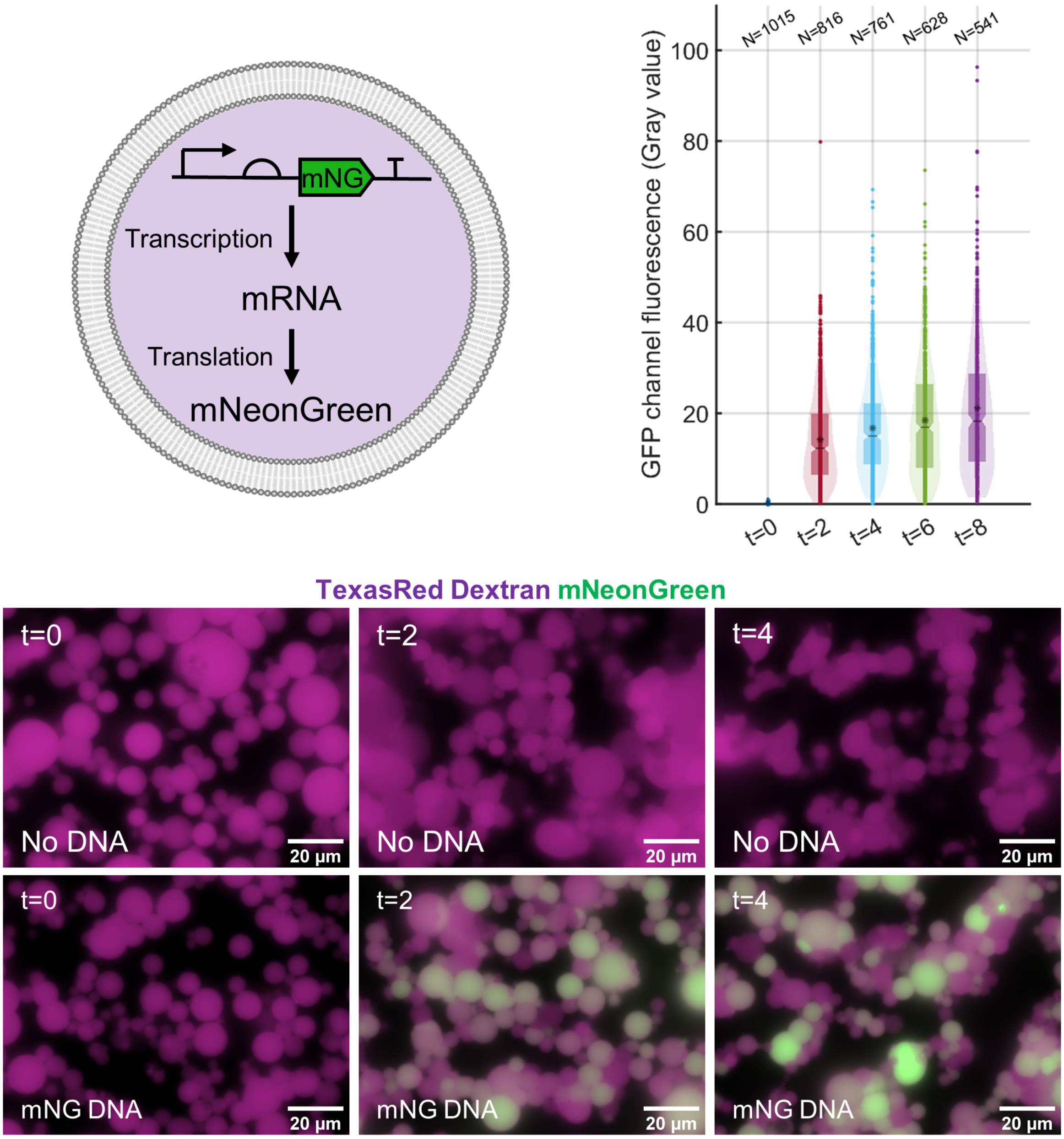


**Figure S3: mNG expression inside GUVs**
Fluorescence microscopy images confirmed mNG was expressed inside GUVs. mNG fluorescence intensity increased over time until components of the CFPS were depleted, and expression began to saturate.

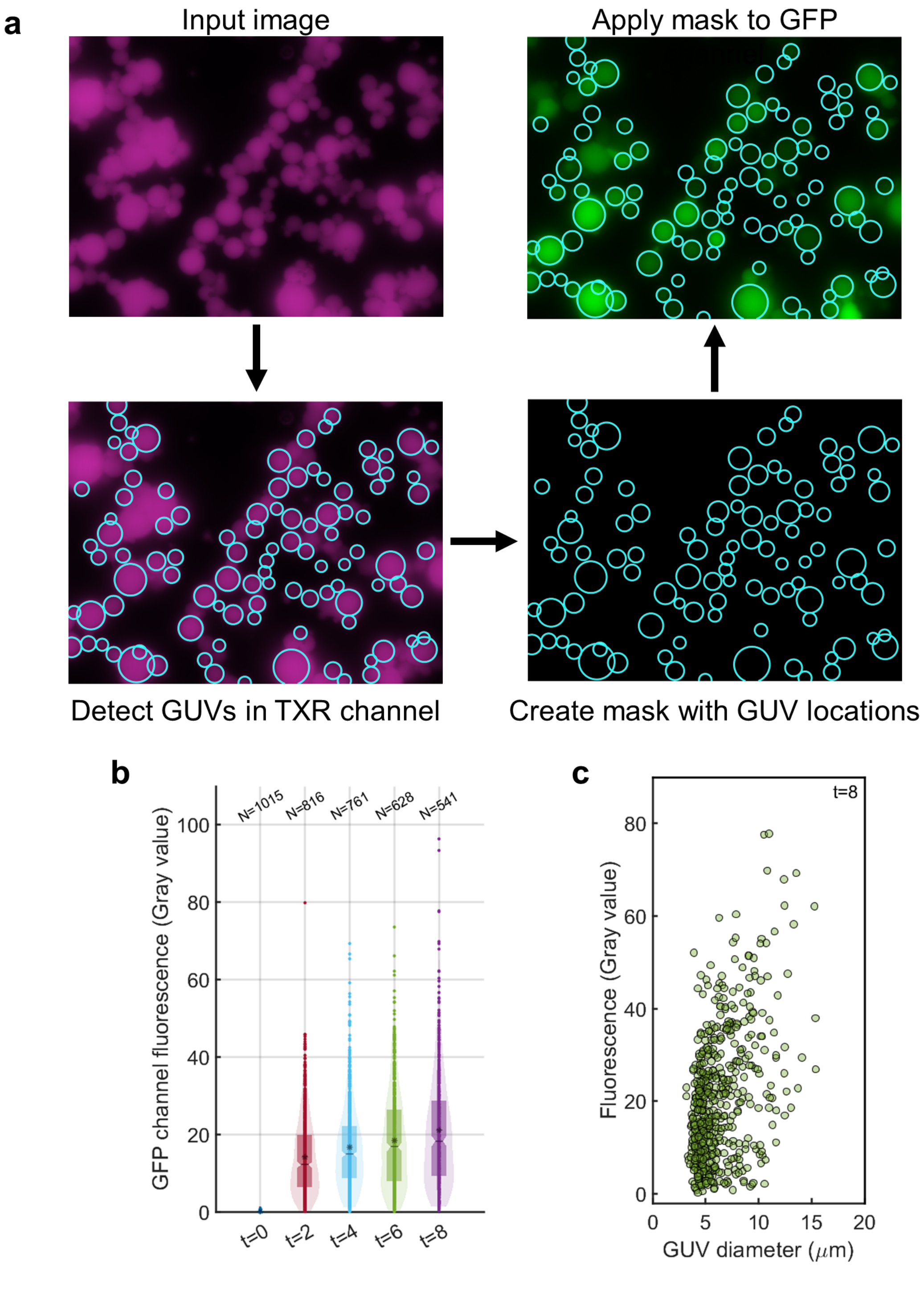


**Figure S4: Quantifying mNG expression inside GUVs**
**a)** TexasRed labelled GUVs present in TXR channel fluorescence microscopy images were used as an input for circular Hough transform. The origin and radii of each GUV were identified and used to create maps. Maps were then applied to the corresponding GFP channel image and mNG expression was determined by extracting the mean pixel intensity inside the individual GUVs. **b)** Image quantification was consistent with the fluorescence microscopy images and illustrated that almost all DNA containing vesicles had a fluorescence value above background (No DNA GUV fluorescence), but mNG expression was highly variable. **c)** Mean pixel intensity (Gray value) vs GUV diameter plots indicated that mNG expression tended to increase with the internal volume of the synthetic cells.

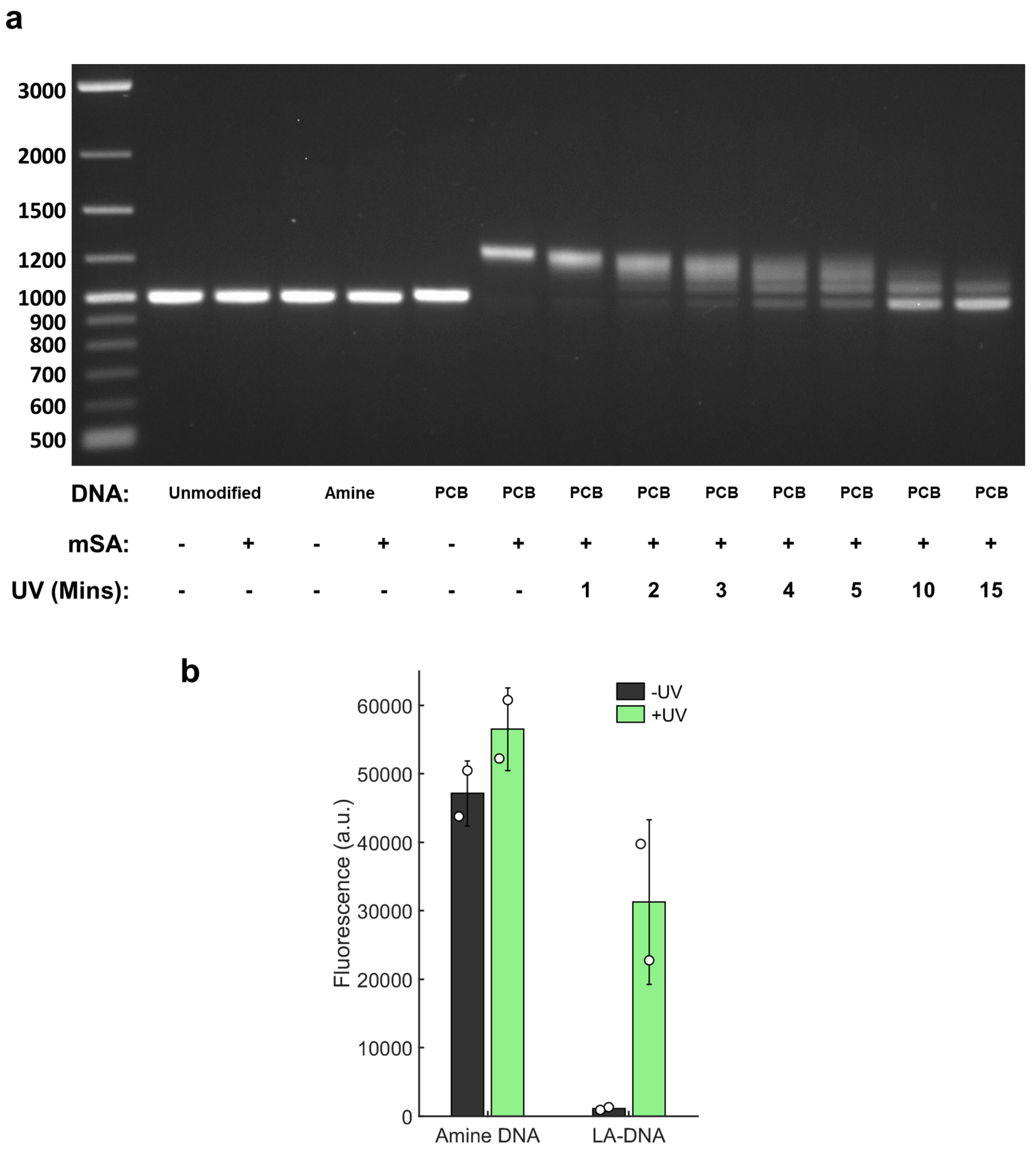


**Figure S5: Characterising light-activated DNA**
**a)** DNA containing PCB groups bound to mSA, forming LA-DNA. LA-DNA had an altering electrophoretic mobility, which caused a band shift compared to PCB DNA. No band shift occurred for unmodified or amine modified DNA, therefore these DNA templates did not bind mSA. mSA was subsequently released from the LA-DNA after UV irradiation. DNA laddering reflects the removal of subsets of the 7 blocking groups. Most blocking groups were removed after 15 mins irradiation. **b)** Minimal mNG was expressed in the absence of UV light, but mNG expression was restored after LA-DNA templates were irradiated with UV light. Samples were irradiated with 0.75 mW UV light for 3 minutes.

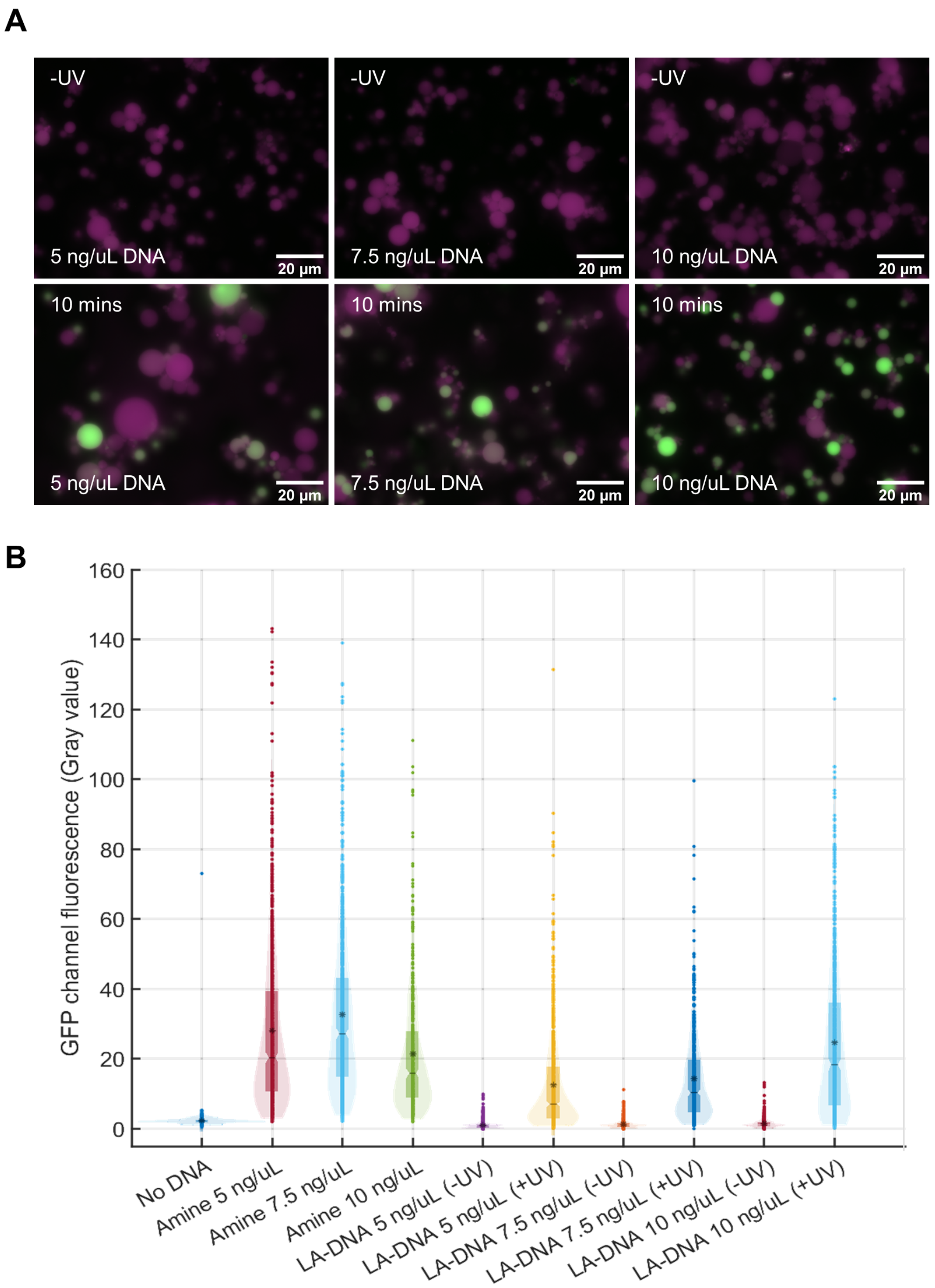


**Figure S6: Light-activated synthetic cells vs DNA template concentration**
**a)** Epifluorescence microscopy images of synthetic cells prepared with 5 ng/$\mu$L, 7.5 ng/$\mu$L or 10 ng/$\mu$L LA-mNG DNA and treated with either no UV light or 0.75 mW UV light for 10 minutes. **b)** Quantification of single vesicle fluorescence for all conditions shown in the microscopy images, compared to the respective DNA concentrations with amine DNA templates. The lower fluorescence output in synthetic cells prepared with 10 ng/$\mu$L DNA was an artificact of the vesicles wetting onto a coverslip that failed to passivate properly.

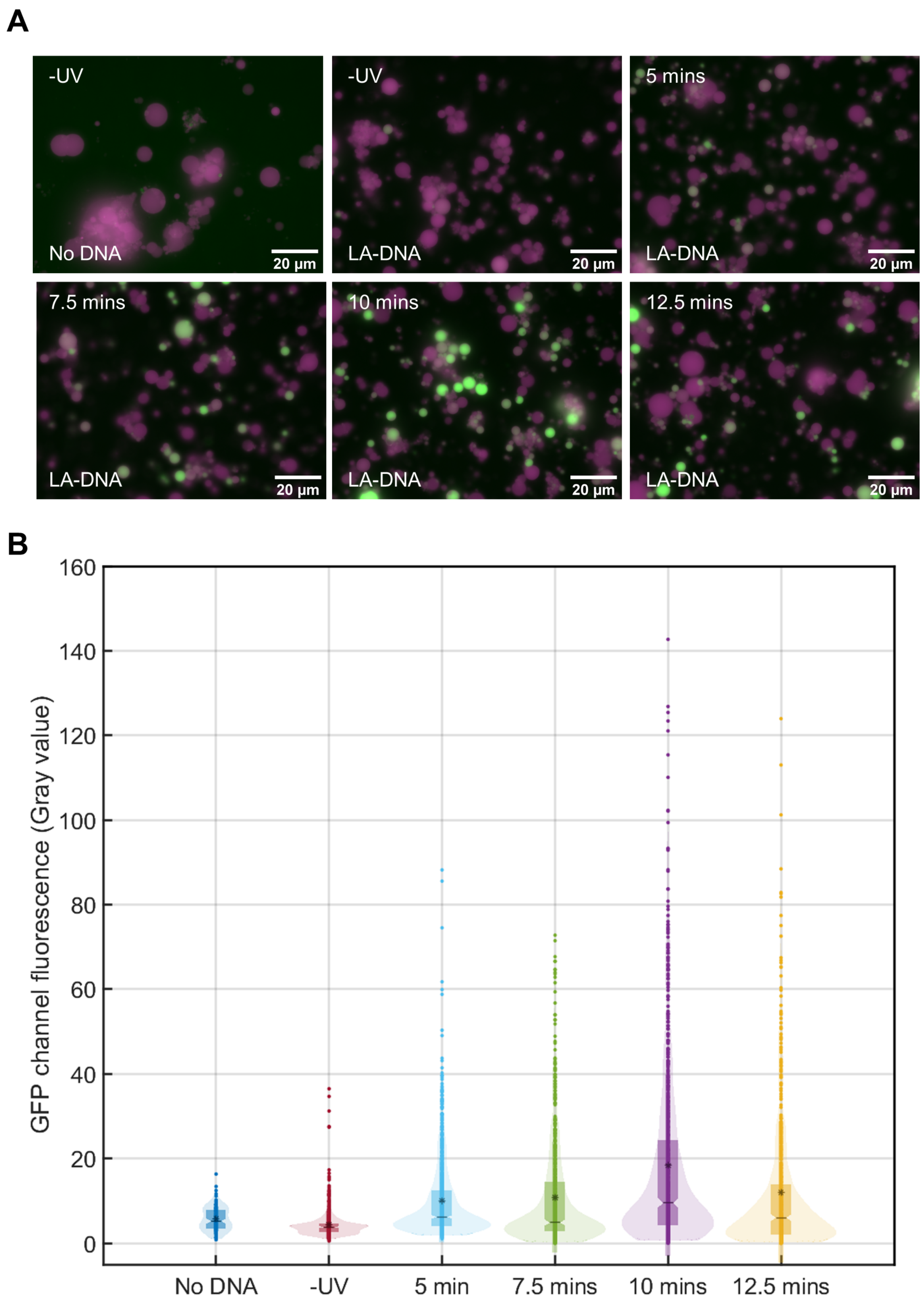


**Figure S7: Light-activated synthetic cells vs UV light exposure time**
**a)** Epifluorescence microscopy images of synthetic cells prepared with 5 ng/$\mu$L LA-mNG DNA and treated with 0.75 mW UV light of increasing duration. **b)** Quantification of single vesicle fluorescence for all conditions shown in the microscopy images, compared to the respective UV light exposure conditions.

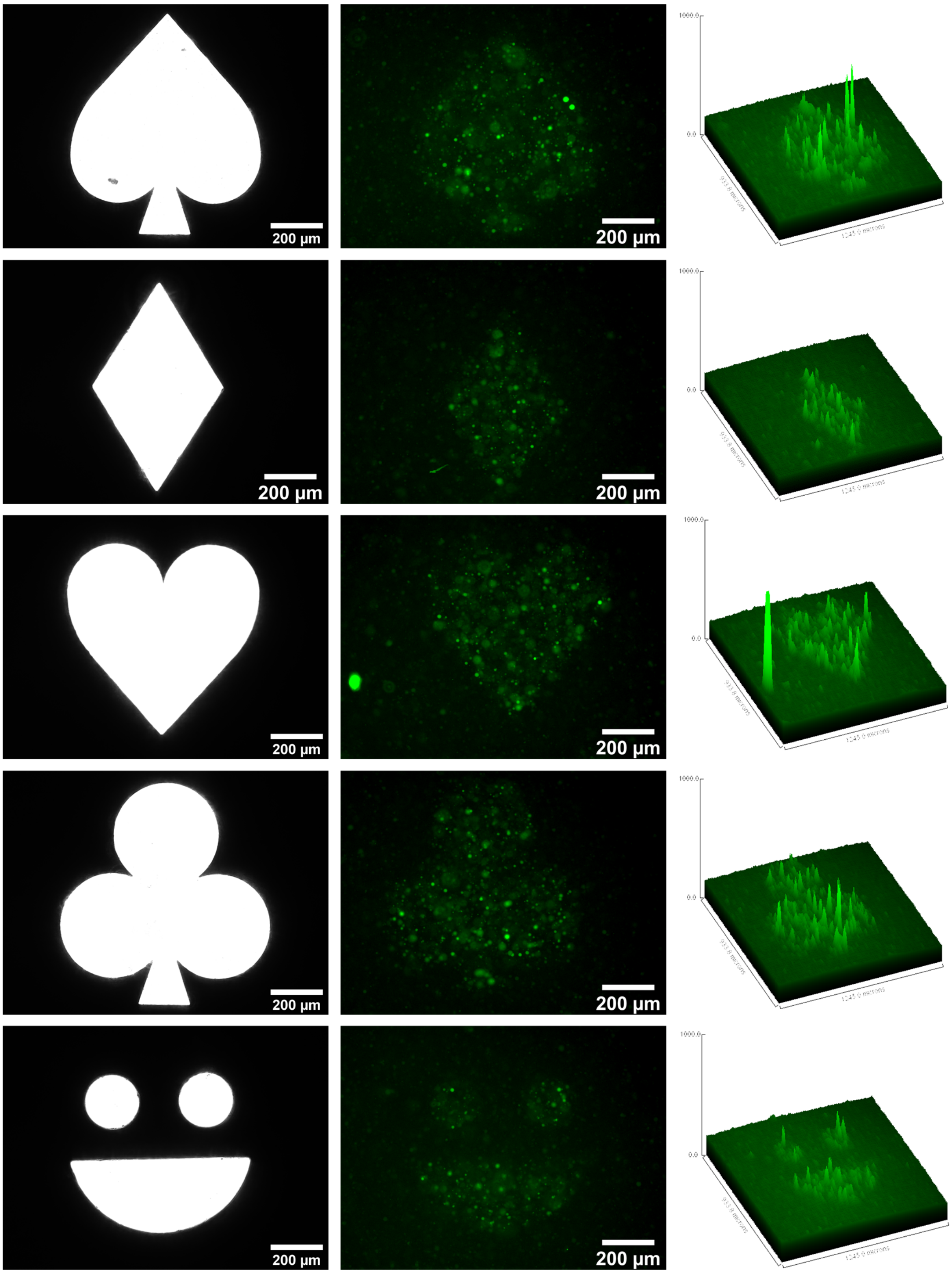


**Figure S8: Patterned LA-SC activation with complex photomasks**
LA-mNG synthetic cells were immobilised in 1.5% ULGP agarose and irradiated with patterned UV light according to photomask designs. mNG was expressed inside GUVs only in the UV exposed areas. Highly patterning fidelity and resolution was demonstrated using the irregular photomask designs that contained features with different dimensions.

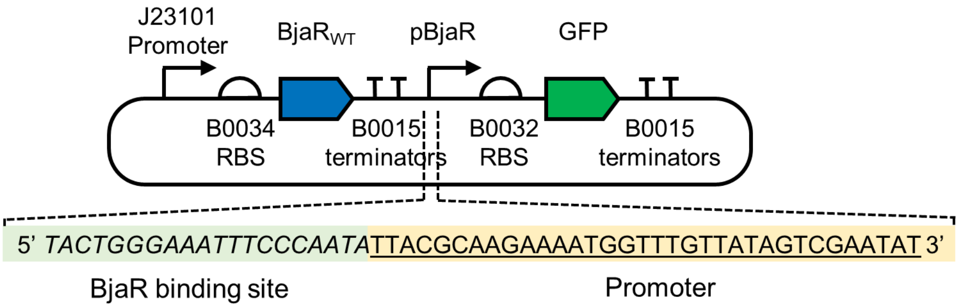


**Figure S9: BjaR reporter plasmids**
pSB1A3 BjaR GFP plasmid. BjaR was expressed under the control of a consituitvely active promoter (J23101). GFP expression was regulated via the pBjaR promoter, and activated upon addition of IV-HSL. The pBjaR promoter comprised the BjaR binding site (Green) and the core pLux promoter sequence (Amber).

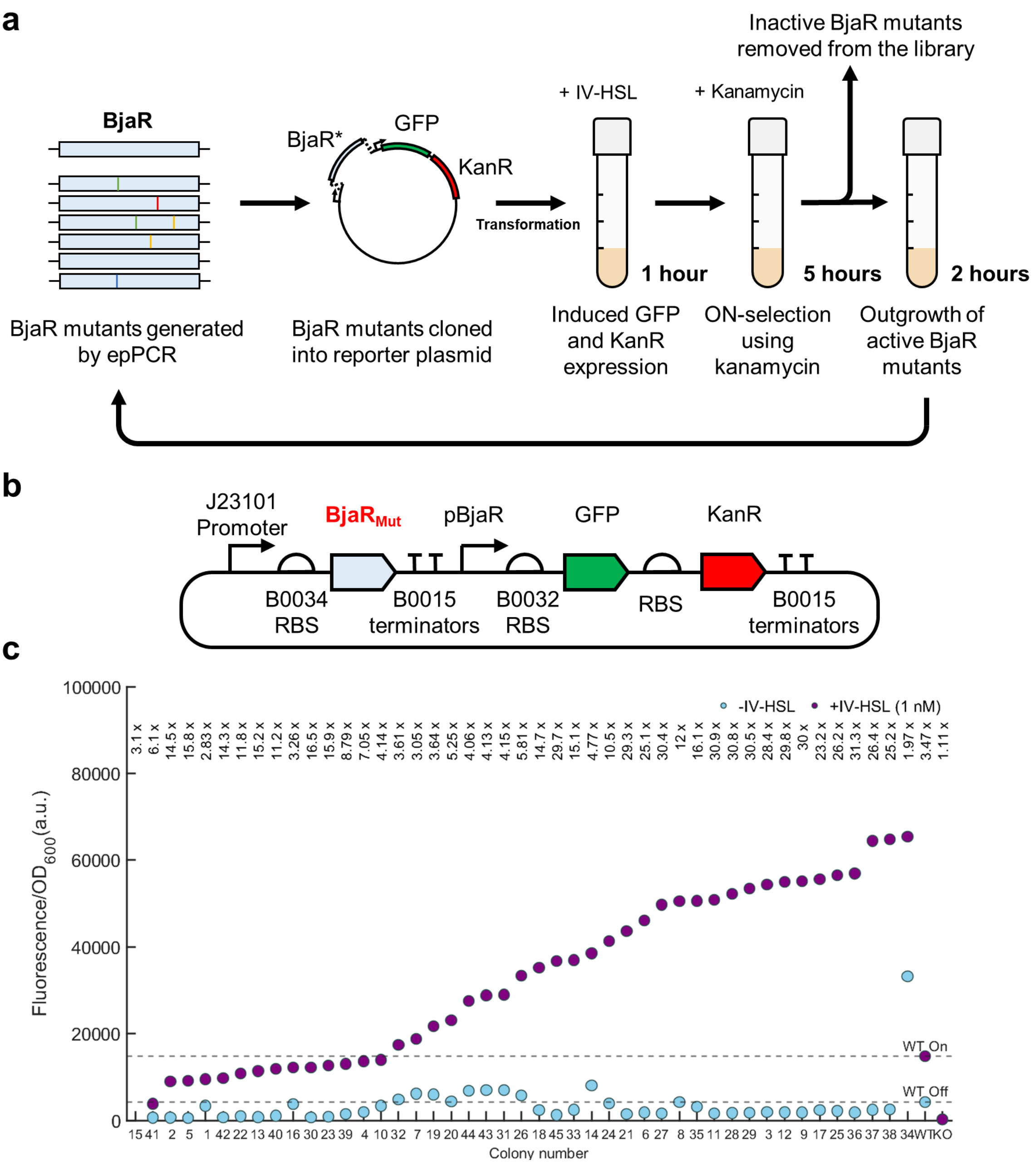


**Figure S10: Screening BjaR mutants from directed evolution**

**a)** The directed evolution workflow used to improve BjaR stringency. **b)** The pSB1A3 BjaR GFP KanR plasmid was constructed to enable activate BjaR mutants to be recovered after ON-selection in Kanamycin. KanR was inserted directed downstream of the GFP gene to create a bicistronic mRNA. Both GFP and KanR expression were activated by the addition of IV-HSL. **c)** GFP expression of 45 colonies containing pSB1A3 $\text{BjaR}_{\text{mut}}$ GFP KanR plasmids were tested in the absence and presence of IV-HSL (1 nM). pSB1A3 $\text{BjaR}_{\text{WT}}$ GFP KanR and pSB1A3 $\text{BjaR}_{\text{KO}}$ GFP KanR were included to enable comparison. Fold change in GFP expression is indicated above the respective colony. Many colonies demonstrated a lower OFF-state and/or improved ON-state.

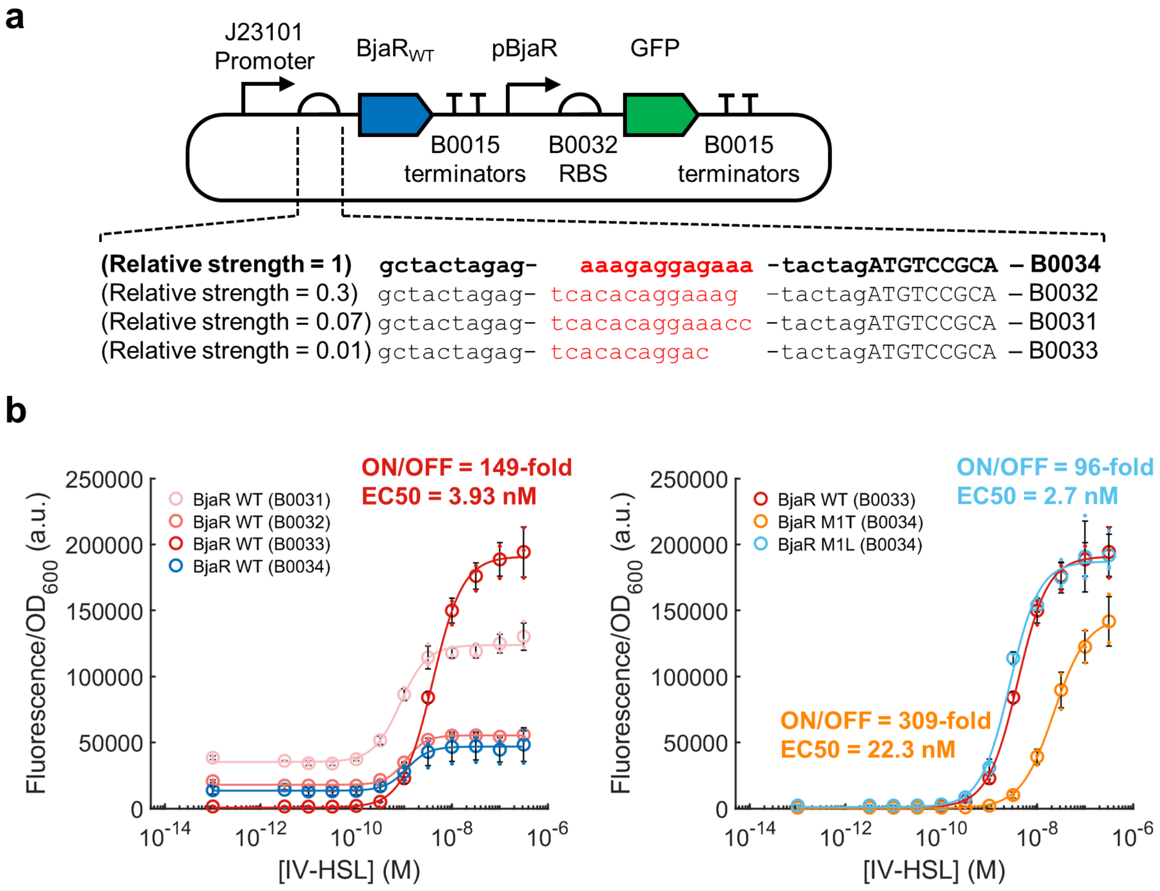


**Figure S11: Light-activated synthetic cells vs DNA template concentration**
**a)** The B0034 RBS in the pSB1A3 $BjaR_{WT}$ GFP plasmid was replaced with RBSs with decreasing relative strengths to decrease BjaR abundance in the cell. **b)** The reporter plasmid containing the weakest RBS (B0033) showed the best dose-response behaviour and closely resembled the influence of the M1L mutation.

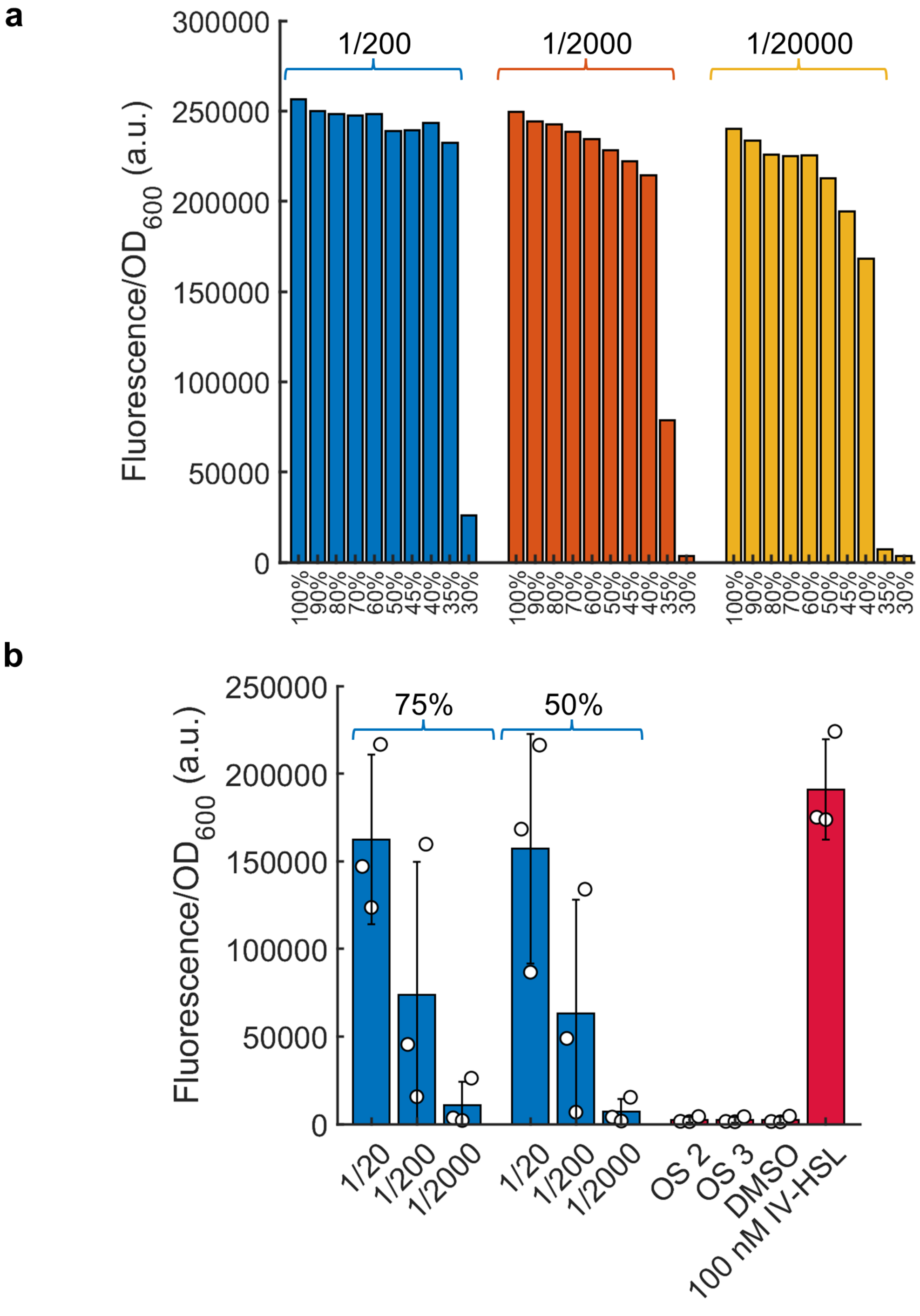


**Figure S12: Light-activated synthetic cells vs DNA template concentration**
**a)** BjaI was expressed in the presence of SAM and IV-CoA in PURExpress at varying dilutions. BjaI expression and IV-HSL synthesis occured even as low as 0.35X the standard PURExpress working concentration. **b)** BjaI-expressing synthetic cells prepared with 0.75X and 0.5X PURExpress produced IV-HSL and activated GFP expression by $\text{BjaR}_{\text{M1L S107R}}$ receiver cells.

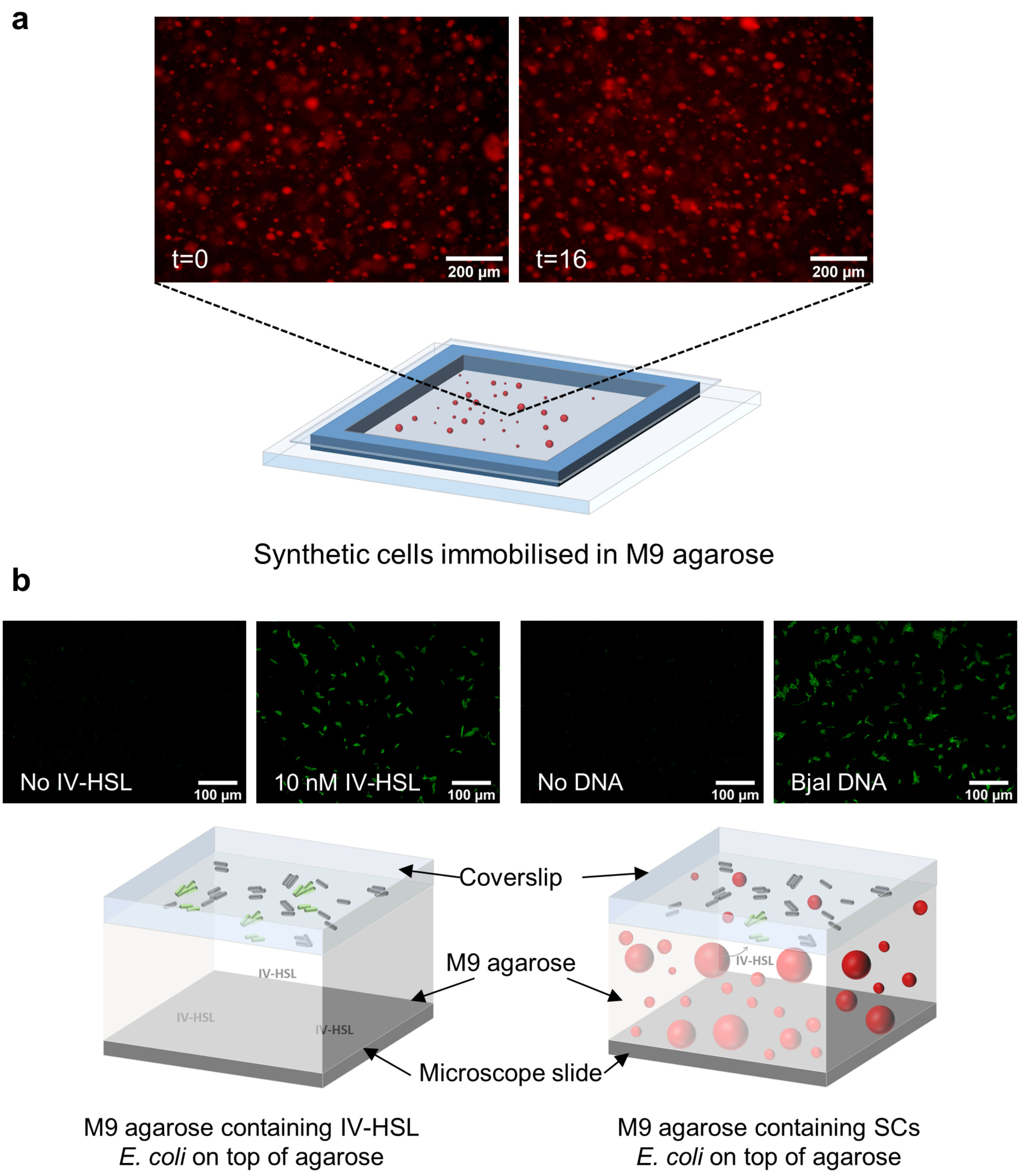


**Figure S13: Synthetic cells in M9 agarose pads**

**a)** Reduced osmolarity synthetic cells prepared with 0.5X PURExpress and 25 $\mu$M TexasRed-Dextran were resuspended in molten 1.5% agarose prepared with M9 minimal media, then set to form agarose pads using gene frames. There was no noticeable increase in background fluorescence nor decrease in the number of synthetic cells present after 16 hours incubation at 37 °C. **b)** BjaR receiver cells were added on top of agarose pads prepared with or without synthetic IV-HSL (10 nM), or pads containing synthetic cells with no DNA template or BjaI DNA. BjaR receiver cells expressed similar amounts of GFP on synthetic IV-HSL pads and BjaI synthetic cell pads confirming the production of IV-HSL and its diffusion across the GUV lipid bilayers and into the agarose.

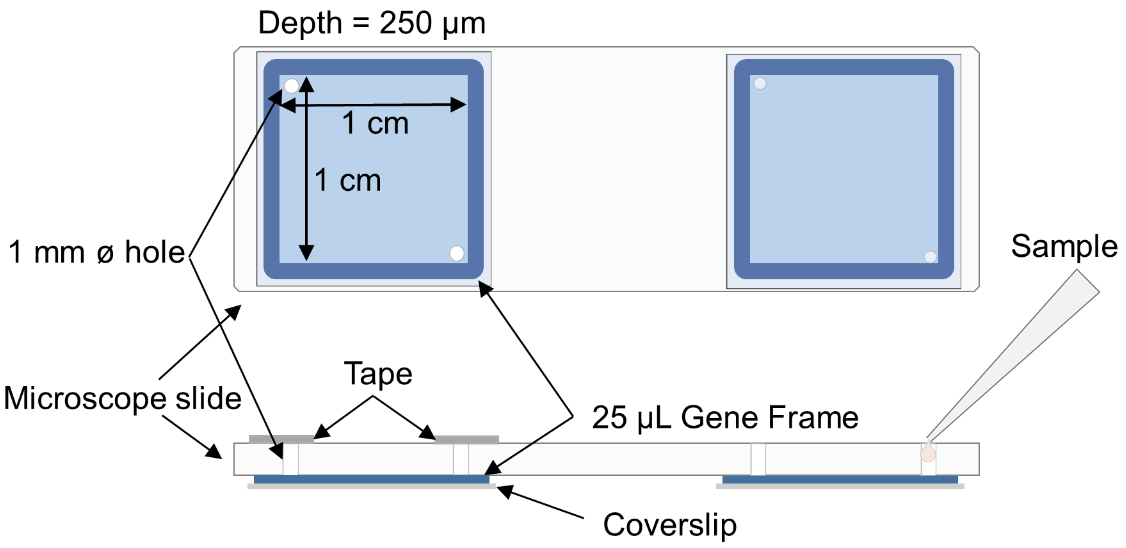


**Figure S14: Imaging chambers**
25 $\mu$L gene frames were placed onto $O_2$ plasma treated 25 mm x 25 mm coverslips, then passivated with BSA. The second side of the gene frame was then sealed onto the drilled coverslips, with opposite corners of the frame aligned with the holes. Sample was introduced through one inlet hole, then both were sealed using double sided tape.

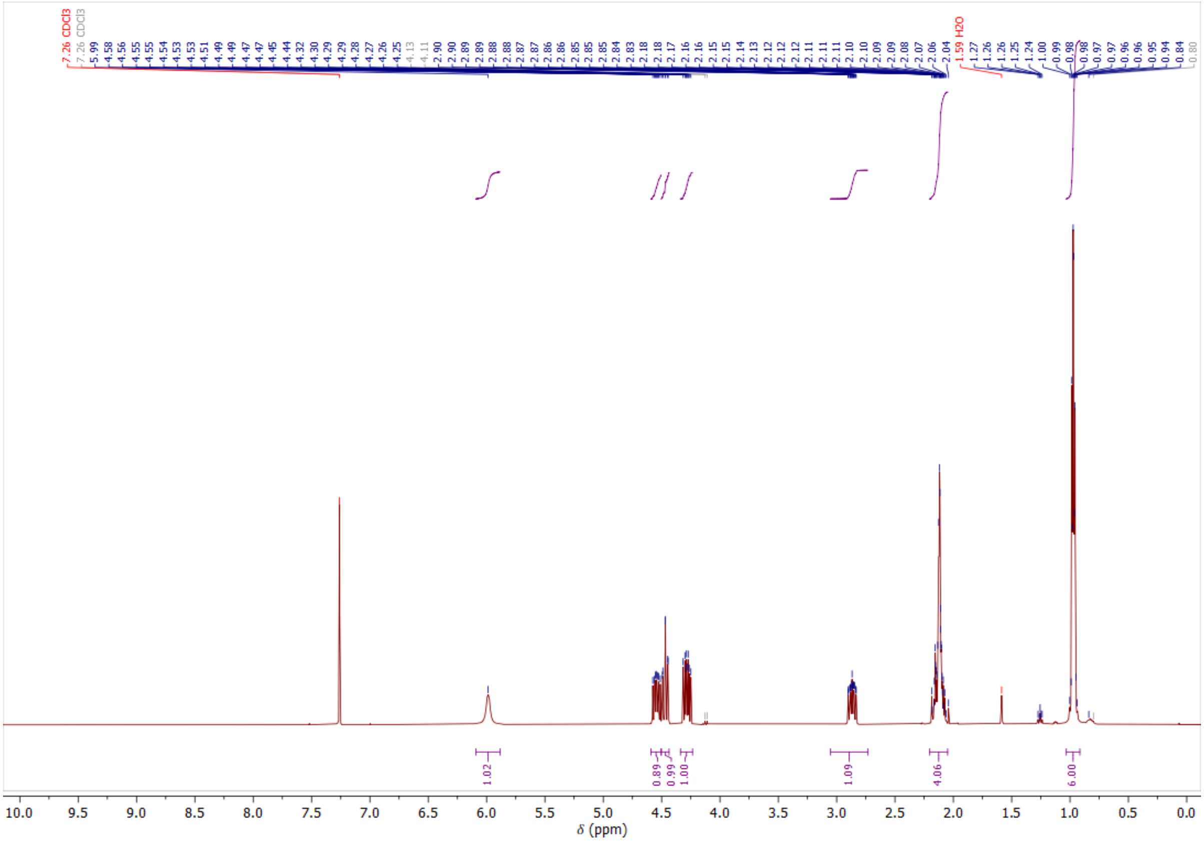


**Figure S15: IV-HSL NMR**
NMR of the synthetic IV-HSL used as a standard in BjaR receiver cell characterisation.

### Supplementary methods

#### pPURE-mNG assembly

The mNG gene was amplified by PCR using Phusion DNA polymerase mastermix containing 1 ng pMAT mNeongreen plasmid and 500 nM T7 for/PURE rev 1 primers with the following thermal cycler programme: 98°C 30 s, 35 cycles of [98 °C for 10 s, 59 °C for 20 s, 72 °C for 45 s], 72 °C for 10 minutes, 4 °C HOLD. 1 $\mu$g of PCR product and PURExpress control template plasmid were digested with XbaI/BamHI-HF for 1 hour at 37 °C and then ran on a 1.2% TAE agarose gel prestained with 3% GelRed at 100 V for 1 hour. Bands corresponding with the double digested DNA were cut from the gel with a scalpel and purified from the gel using QIAquick gel extraction purification kit. The mNG gene insert was ligated with 50 ng plasmid backbone using T4 DNA ligase (3:1 insert:vector ratio) and 2 $\mu$L of ligation was transformed into 10 $\mu$L *E. coli* XL10 Gold cells by heat shock. Cells were incubated in 500 $\mu$L SOC media for 45 minutes, then 100 $\mu$L was plated onto LB agar containing 100 $\mu$g/mL ampicillin. Plates were incubated at 37°C for ∼14 hours. Colonies were picked with pipette tips and grown as 5 mL liquid cultures in LB + ampicillin (100 ug/mL) at 37 °C, 225 rpm, overnight. Plasmids were harvested from 3 mL overnight culture using a QIAprep spin mini-prep kit and verified by Sanger sequencing.

#### pPURE-mNG T7g10 assembly

The T7g10 leader sequence was introduced into the pPURE-mNG plasmid via the recombination of a single DNA fragment with homologous ends, denoted mNG T7g10 BB [42]. mNG T7g10 BB was prepared by nested PCR. First, mNG T7g10 fragment 1 was amplified with Phusion DNA polymerase mastermix (2X) using 500 nM mNG_T7g10_for1 and mNG_T7g10_rev1 primers and 1 ng pPURE-mNG template with the following thermal cycler programme: 98 °C for 30s, 35 cycles of [98 °C for 10 s, 63 °C for 20 s, 72 °C for 1 min 30 s], 72 °C for 10 mins, 4 °C HOLD. mNG T7g10 BB was then amplified from 1 ng mNG T7g10 fragment 1 using Phusion DNA polymerase mastermix (2X) and 500 nM mNG_T7g10_for2 and mNG_T7g10_rev2 primers with the following thermal cycler programme: 98 °C for 30s, 35 cycles of [98 °C for 10 s, 63 °C for 20 s, 72 °C for 1 min 30 s], 72 °C for 10 mins, 4 °C. After amplification, 10 U of DpnI was added to the reaction mix and incubated at 37 °C for 1 hour, followed by 20 mins at 80 °C. The PCR product was purified from the reaction mixture using a QIAquick spin PCR purification kit. 100 ng of mNG T7g10 BB was transformed into 10 $\mu$L *E. coli* XL10-Gold cells by heat shock. Cells were incubated in 500 $\mu$L SOC media for 45 minutes, then 100 $\mu$L was plated onto LB agar containing 100 $\mu$g/mL ampicillin. Plates were incubated at 37 °C for ∼14 hours. Colonies were picked with pipette tips and grown as 5 mL liquid cultures in LB + ampicillin (100 $\mu$g/mL) at 37 °C, 225 rpm, overnight. Plasmids were harvested from 3 mL overnight culture using a QIAprep spin mini-prep kit and verified by Sanger sequencing.

#### pPURE-mNG g10 assembly

The 10 AA leader sequence (9 AAs from the T7 bacteriophage major capsid protein + 1 additional AA (Histidine) present in [28]) was introduced into the pPURE-mNG plasmid via the recombination of 2 DNA strands with homologous ends, denoted mNG g10 insert and mNG g10 BB. mNG g10 insert was amplified with Phusion DNA polymerase mastermix (2X) using 500 nM mNG_g10_for and PURE_rev2 primers and 1 ng pPURE-mNG T7g10 template with the following thermal cycler programme: 98 °C for 30 s, 35 cycles of [98 °C for 10 s, 66 °C for 20 s and 72 for °C for 30 s], 72 °C for 5 mins, 4 °C HOLD. mNG g10 BB was amplified with Phusion DNA polymerase mastermix (2X) using 500 nM PURE_for1 and mNG_g10_rev primers and 1 ng pPURE-mNG template with the following thermal cycler programme: 98 °C 30 s, 35 cycles of [98 °C for 10 s, °C for 20 s and 72 °C for 1 min 15 s], 72 °C for 5 mins, 4 °C HOLD. After amplification, 10 U of DpnI was added to the reactions and incubated at 37 °C for 1 hour, followed by 20 mins at 80 °C. The PCR products were purified from the reaction mixture using a QIAquick spin PCR purification kit. Insert and BB were combined at a 3:1 mol ratio (100 ng of mNG g10 BB) and transformed into 10 $\mu$L *E. coli* XL10-Gold cells by heat shock. Cells were incubated in 500 $\mu$L SOC media for 45 minutes, then 100 $\mu$L was plated onto LB agar containing 100 $\mu$g/mL ampicillin. Plates were incubated at 37 °C for ~14 hours. Colonies were picked with pipette tips and grown as 5 mL liquid cultures in LB + ampicillin (100 $\mu$g/mL) at 37 °C, 225 rpm, overnight. Plasmids were harvested from 3 mL overnight culture using a QIAprep spin mini-prep kit and verified by Sanger sequencing.

#### pPURE-mVenus T7g10 assembly

The T7g10 leader sequence was introduced into the pPURE-mVenus plasmid via the recombination of 2 DNA strands with homologous ends, denoted mVenus_T7g10_insert and mVenus_T7g10_BB. mVenus_T7g10_insert was amplified with Phusion DNA polymerase mastermix (2X) using 500 nM mV_T7g10_for and PURE_rev2 primers and 1 ng pPURE-mVenus template with the following thermal cycler programme: 98 °C for 30 s, 35 cycles of [98 °C for 10 s, 66 °C for 20 s and 72 for °C for 30 s], 72 °C for 10 mins, 4 °C HOLD. mVenus T7g10 BB was amplified with Phusion DNA polymerase mastermix (2X) using 500 nM PURE_for1 and mV_T7g10_rev primers and 1 ng pPURE-T7g10 mNG template with the following thermal cycler programme: 98 °C 30 s, 35 cycles of [98 °C for 10 s, 66 °C for 20 s and 72 °C for 1 min 15 s], 72 °C for 5 mins, 4 °C HOLD. Two fragment homologous recombination was performed as described above.

#### pPURE-BjaI assembly

The BjaI gene sequence was obtained from the European Nucleotide Archive database (Sequence BA000040.2)[25]. Short ~60 nt oligonucleotides (BjaI_for1-9/Bjai_rev1-8) with ~20 nt terminal overhangs were designed to collectively encode the full BjaI gene sequence. Oligos BjaI_for1-9 and BjaI_rev1-8 were pooled to a final concentration of 5 $\mu$M, then annealed and elongated via polymerase chain assembly [43] (100 nM oligo mix, 1X Phusion DNA polymerase mastermix, 3% DMSO) with the following thermal cycler programme (98 °C 2 mins, 30 cycles of [98 °C for 10 s, 60 °C for 20 s and 72 °C for 15 s], 72°C for 5 mins, 4 °C HOLD). The complete BjaI gene was amplified from the pool of assembled oligos via a second PCR step with amplification primers BjaI_amp_for and BjaI_amp_rev. BjaI PCR product and pPURE-mNG plasmid were double digested with XbaI/BamHI-HF and NdeI/EcoRI-HF and ran on a 1.2% TAE agarose gel. Digested DNA was cut out from the agarose gel and purified using the Monarch gel extraction purification kit. The BjaI gene was ligated into digested pPURE plasmid using the same protocol as described above.

#### pSB1A3 BjaR$_{\mathrm{KO}}$ GFP assembly

pSB1A3 BjaR$_{\mathrm{KO}}$ GFP BB was prepared by the homologous recombination of a single DNA fragment with homologous ends, denoted BjaR$_{\mathrm{KO}}$ BB. To knockout BjaR activity, the BjaR gene was truncated by introducing two stop codons followed by a randomised DNA sequence after position 12. Nucleotides for AA1-12 were left to assist in cloning the plasmid. BjaR$_{\mathrm{KO}}$ BB was amplified with Phusion DNA polymerase mastermix (2X) using 500 nM BjaR_KO_for and BjaR_KO_rev primers and 1 ng pSB1A3 BjaR$_{\mathrm{WT}}$ GFP template with the following thermal cycler programme: 98 °C for 30s, 35 cycles of [98 °C for 10 s, 65 °C for 20 s, 72 °C for 1 min], 72 °C for 5 mins, 4 °C HOLD. Single fragment homologous recombination was performed as described above.

#### pSB1A3 BjaR$_{\mathrm{WT}}$ GFP KanR and pSB1A3 BjaR$_{\mathrm{KO}}$ GFP KanR assembly

pSB1A3 BjaR$_{\mathrm{KO/WT}}$ GFP KanR were prepared via the recombination of two DNA strands with homologous ends, denoted BjaR$_{\mathrm{KO/WT}}$ KanR BB and KanR insert. The BjaR$_{\mathrm{KO/WT}}$ KanR BB fragments were amplified with Phusion DNA polymerase mastermix (2X) using 500 nM pSB1A3_for3 and pSB1A3_rev3 primers and 1 ng pSB1A3 BjaR$_{\mathrm{KO/WT}}$ GFP templates with the following thermal cycler programme: 98 °C for 30 s, 35 cycles of [98 °C for 10 s, 65 °C/63 °C for 20 s and 72 °C for 1 min/2 mins], 72 °C for 10 mins, 4 °C HOLD. The KanR insert fragment was created by nested PCR. KanR was first amplified with Phusion DNA polymerase mastermix (2X) using 500 nM KanR_for1 and KanR_rev primers and 1 ng pET24a template with the following thermal cycler programme: 98 °C for 30 s, 35 cycles of [98 °C for 10 s, 66 °C for 20 s and 72 °C for 30 s], 72 °C for 5 mins, 4 °C HOLD. This PCR product was purified using a QIAquick PCR purification kit then 1 ng was used as the DNA template in a second PCR with 500 nM KanR_for2 and KanR rev primers according to the following protocol: 98 °C for 30 s, 35 cycles of [98 °C for 10 s, 65 °C for 20 s and 72 °C for 30 s], 72 °C for 5 mins, 4 °C HOLD. Two fragment homologous recombination was performed as described above.

#### pSB1A3 BjaR$_{\mathrm{M1L\ S107R}}$ GFP and pSB1A3 BjaR$_{\mathrm{M1T}}$ GFP assembly

pSB1A3 BjaR$_{\mathrm{M1T/M1L\ S107R}}$ GFP were prepared by transferring the BjaR$_{\mathrm{M1T/M1L\ S107R}}$ genes from the KanR containing reporter plasmid backbone used in directed evolution workflow back into the KanR-free backbone. pSB1A3 BjaR$_{\mathrm{M1T/M1L\ S107R}}$ GFP were prepared via the recombination of two DNA strands with homologous ends, denoted pSB1A3 GFP BB and BjaR$_{\mathrm{M1T/M1L\ S107R}}$ insert. The pSB1A3 GFP BB fragment was amplified with Phusion DNA polymerase mastermix (2X) using 500 nM pSB1A3_for2 and pSB1A3_rev2 primers and 1 ng pSB1A3 BjaR$_{\mathrm{WT}}$ GFP template with the following thermal cycler programme: 98 °C for 30 s, 35 cycles of [98 °C for 10 s, 63 °C for 20 s and 72 °C for 2 mins], 72 °C for 10 mins, 4 °C HOLD. The BjaR$_{\mathrm{M1T/M1L\ S107R}}$ insert fragments were amplified with Phusion DNA polymerase mastermix (2X) using 500 nM pSB1A3_for1 and pSB1A3_rev1 primers and 1 ng pSB1A3 BjaR$_{\mathrm{M1T/M1L\ S107R}}$ GFP KanR templates with the following thermal cycler programme: 98 °C for 30 s, 35 cycles of [98 °C for 10 s, 63 °C for 20 s and 72 °C for 30 s], 72 °C for 5 mins, 4 °C HOLD. Two fragment homologous recombination was performed as described above.

#### pSB1A3 (B0031-33) BjaR$_{\mathrm{WT}}$ GFP assembly

The Bba_B0034 RBS from pSB1A3 BjaR$_{\mathrm{WT}}$ GFP was replaced by weaker Bba_B0031/32/33 RBS' via the recombination of 2 DNA strands with homologous ends, denoted pSB1A3 BjaR$_{\mathrm{WT}}$ B0031-33 GFP BB and B0031/B0032/B0033 insert. The pSB1A3 BjaR$_{\mathrm{WT}}$ B0031/B0032/B0033 GFP BB fragments amplified with Phusion DNA polymerase mastermix (2X) using 500 nM pSB1A3_for1 and B0031-33_rev primers and 1 ng pSB1A3 BjaR$_{\mathrm{WT}}$ GFP template with the following thermal cycler programme: 98 °C for 30 s, 35 cycles of [98 °C for 10 s, 66 °C for 20 s and 72 °C for 1 min 15 s], 72 °C for 10 mins, 4 °C HOLD. The B0031/B0032/B0033 insert fragments were amplified with Phusion DNA polymerase mastermix (2X) using 500 nM B0031/B0032/B0033_for and pSB1A3_rev1 primers and 1 ng pSB1A3 BjaR$_{\mathrm{WT}}$ GFP template with the following thermal cycler programme: 98 °C for 30 s, 35 cycles of [98 °C for 10 s, 66 °C for 20 s and 72 °C for 30 s], 72 °C for 5 mins, 4 °C HOLD. 2 fragment homologous recombination was performed as described previously.

#### pSB1A3 BjaR$_{\mathrm{M1L}}$ GFP assembly

The M1L mutation was introduced into pSB1A3 BjaR$_{\mathrm{WT}}$ GFP via the recombination of 2 DNA strands with homologous ends, denoted pSB1A3 BjaR$_{\mathrm{M1L}}$ GFP BB and BjaR$_{\mathrm{M1L}}$ insert. pSB1A3 BjaR$_{\mathrm{M1L}}$ GFP BB was amplified with Phusion DNA polymerase mastermix (2X) using 500 nM pSB1A3_for2 and pSB1A3_rev2_M1L primers and 1 ng pSB1A3 BjaR$_{\mathrm{WT}}$ GFP template with the following thermal cycler programme: 98 °C for 30 s, 35 cycles of [98 °C for 10 s, 66 °C for 20 s and 72 °C for 1 min 30 s], 72 °C for 10 mins, 4 °C HOLD. BjaR$_{\mathrm{M1L}}$ insert was amplified with Phusion DNA polymerase mastermix (2X) using 500 nM pSB1A3_for1_M1L and pSB1A3_rev1 and 1 ng pSB1A3 BjaR$_{\mathrm{WT}}$ GFP template with the following thermal cycler programme: 98 °C for 30 s, 35 cycles of [98 °C for 10 s, 66 °C for 20 s and 72 °C for 15 s], 72 °C for 5 mins, 4 °C HOLD. Two fragment homologous recombination was performed as described previously.

#### pSB1A3 (B0033) BjaR$_{S107R}$ GFP assembly

pSB1A3 BjaR$_{S107R}$ (B0033) GFP was prepared by introducing the S107R mutation into the BjaR gene within the B0033 backbone. pSB1A3 BjaR$_{S107R}$ (B0033) GFP was prepared via the recombination of two DNA strands with homologous ends, denoted BjaR$_{S107R}$ (B0033) BB and BjaR$_{S107R}$ insert. BjaR$_{S107R}$ (B0033) BB was amplified with Phusion DNA polymerase mastermix (2X) using 500 nM pSB1A3_for2 and S107R_rev primers and 1 ng pSB1A3 BjaR$_{WT}$ (B0033) GFP template with the following thermal cycler programme: 98 °C for 30 s, 35 cycles of [98 °C for 10 s, 63 °C for 20 s and 72 °C for 1 min 30 s], 72 °C for 10 mins, 4 °C HOLD. BjaR$_{S107R}$ insert was amplified with Phusion DNA polymerase mastermix (2X) using 500 nM S107R_for and pSB1A3_rev1 primers and 1 ng pSB1A3 BjaR$_{WT}$ (B0033) GFP templates with the following thermal cycler programme: 98 °C for 30 s, 35 cycles of [98 °C for 10 s, 63 °C for 20 s and 72 °C for 15 s], 72 °C for 5 mins, 4 °C HOLD. Two fragment homologous recombination was performed as described above.

| Name | Sequence (5'-3') |
|---|---|
| T7_For | GAAATTAATACGACTCACTATAGGGTCTAG |
| T7_For_amine | GAAAT**T**AA**T**ACGAC**T**CAC**T**A**T**AGGG**T**C**T**AG |
| T7_For_PCB | GAAATTAATACGACTCACTATAGGGTCTAG |
| PURE_rev1 | GATATAGTTCCTCCTTTCAG |
| PURE_rev2 | GCCTCCTGCAGGTTAACCTTAC |
| PURE_for1 | CTCGAGTAAGGTTAACCTGCAGGAG |
| mNG_T7g10_for1 | AGATATACCATGGCTAGCATGACTGGTGGACAGCAACA TATGGTGAGCAAAGGCGAAGAG |
| mNG_T7g10_rev1 | TATTTCTAGAGGGAAACCGTTGTGGTCTCCCTAGACCC TATAGTGAGTCGTATTAATTTC |
| mNG_T7g10_for2 | CTCTAGAAATAATTTTGTTTAACTTTAAGAAGGAGATA TACCATGGCTAGCATGACTG |
| mNG_T7g10_rev2 | GCCATGGTATATCTCCTTCTTAAAGTTAAACAAAATTA TTTCTAGAGGGAAACCGTTGTG |
| mNG_g10_for | ATGGCTAGCATGACTGGTGGAC |
| mNG_g10_rev | GTCCACCAGTCATGCTAGCCATATGTATACCTCCTTCT TAAAGTTAAAC |
| mV_T7g10_for | CATGACTGGTGGACAGCAACATATGGTGAGCAAGGGC GAGGAGC |
| mV_T7g10_rev | CATATGTTGCTGTCCACCAGTCATGC |

| | |
|---|---|
| BjaI_for1 | GAAGGAGGTATACATATGATTCACGCAATTTCCGCGGTCAATCGCCACTTATACGAGGAC |
| BjaI_for2 | GCGTCATGACATCTTTGTCGAGGAGCGGCACTGGGAGACGCTGCGCAGGCCGGATGGCCG |
| BjaI_for3 | GAGGATACCGTCTATCTGCTTGCGCTGGAGGGACGGCGCGTCGTCGGCGGCCACCGGCTC |
| BjaI_for4 | CTCGATGATGAGCGAGGTCTTCCCGCATCTGGCGGCGGTTCGCGGCTGCCCCTCGGATC |
| BjaI_for5 | CTACTTCGTCGTCCGCGATCGCCGCGACGGCGCGCTCAACCTGCAACTGATGGCGGCG |
| BjaI_for6 | GAATCGCGCAGGTCAGCGCGATCATGGAAACCTGGTGGTTGCCGCGCTTCCACGAGGCCG |
| BjaI_for7 | CTGCCGGCTCTGGTCGAGAACGCCTGGACCATGGCGGCCACCGTCGACATTCGTCGCCAG |
| BjaI_for8 | GATCGCATCGGCATGCCTTCCATCGTGCAACAGGACGGCAGGACGGCCCGCGTCTGGACGCCGTCGC |
| BjaI_for9 | GCCGCGCAACGAAAGAGCGCCTGATGAAATGGATCCCGGGAATTCTCGA |
| BjaI_rev1 | GCTCTTTCGTTGCGCGGCAGCGAGGCCGCAGCACGGGCGACGGCGTCCAGACG |
| BjaI_rev2 | GAAGGCATGCCGATGCGATCATGCAGGACATCGAGCGTCTGGCGACGAATGTCGACG |
| BjaI_rev3 | CTCGACCAGAGCCGGCAGGCCGAGCGGCGTCACGACGAAGCCGGCCTCGTGGAAGCGCG |
| BjaI_rev4 | GCTGACCTGCGCGATTCCCTGGTCGAGGCAGAACTCCTGCACCGCCGCCATCAGTTGCAG |
| BjaI_rev5 | GATCGCGGACGACGAAGTAGCGCGACCATTCCCAGATCAGCGGATCCGAGGGGCAGCCGC |
| BjaI_rev6 | GACCTCGCTCATCATCGAGGGCTTGGTCGTGGGGTAGAGCCGGTGGCCGCCGAC |
| BjaI_rev7 | GCAGATAGACGGTATCCTCGTCGTCATAGGAATCGACCTCGCGGCCATCCGGCCTGCGC |
| BjaI_rev8 | GACAAAGATGTCATGACGCAGCCGGAAATGCTGCTCGAGTACGTCCTCGTATAAGTGGCG |
| BjaI_for_amp | GAAGGAGGTATACATATGATTCACGC |
| BjaI_rev_amp | CGAGAATTCCCGGGATCCATTTC |
| BjaR_KO_For | TGACATTCTCTCAAAGTATTATGCAGGGCCATCCGTCACAAAATTATCAAC |
| BjaR_KO_Rev | CCTGCATAATACTTTGAGAGAATGTCACAGGGCTTCACGCCCATAATCTAC |

| | |
|---|---|
| KanR_for1 | AGTAAATCTAAGCAGGTCCGCATGAGCCATATTCAACG GGAAAC |
| KanR_for2 | CATGGCATGGATGAACTATACAAATAATAAAGTAAATC TAAGCAGGTCCGC |
| KanR_rev | CTGACCGATAAGCCGGCTCTAGTATTATTAG AAAAACTCATCGAGCATC |
| pSB1A3_for1 | GCTACTAGAGAAAGAGGAGAAATACTAG |
| pSB1A3_for2 | TACTAGAGCCAGGCATCAAATAAAACG |
| pSB1A3_for3 | TACTAGAGCCGGCTTATCGGTCAG |
| pSB1A3_rev1 | CGTTTTATTTGATGCCTGGCTCTAGTA |
| pSB1A3_rev2 | CTAGTATTTCTCCTCTTTCTCTAGTAGC |
| pSB1A3_rev3 | TTATTATTTGTATAGTTCATCCATGCCATG |
| pSB1A3_for1_M1L | GCTACTAGAGAAAGAGGAGAAATACTAGCTG |
| pSB1A3_rev2_M1L | CAGCTAGTATTTCTCCTCTTTCTCTAGTAGC |
| B0030_for | CTAGAGATTAAAGAGGAGAAATACTAGATGTCCGCAGT AGATTATGGGC |
| B0030_rev | GTATTTCTCCTCTTTAATCTCTAGTAGCTAGCACTGTA CCTAGGAC |
| B0031_for | CTAGAGTCACACAGGAAACCTACTAGATGTCCGCAGTA GATTATGGGC |
| B0031_rev | GTAGGTTTCCTGTGTGACTCTAGTAGCTAGCACTGTAC CTAGGAC |
| B0032_for | CTAGAGTCACACAGGAAAGTACTAGATGTCCGCAGTAG ATTATGGGC |
| B0032_rev | CTAGTACTTTCCTGTGTGACTCTAGTAGCTAGCACTGT ACCTAGGAC |
| B0033_for | CTACTAGAGTCACACAGGACTACTAGATGTCCGCAGTA GATTATGGGC |
| B0033_rev | GTAGTCCTGTGTGACTCTAGTAGCTAGCACTGTACCTA GGAC |
| S107R_for | GAATTAGAACCACGCGCCGC |
| S107R_rev | GCGGCGCGTGGTTCTAATTC |

**Table 1: Primer sequences**
Bold bases indicate a C6-thymine modified base. Underlined bases indicate a PCB-modified thymine base.

167
